# Bead-based RNA Detection with Cas13a: Multi-scale Design and Validation

**DOI:** 10.64898/2026.07.27.739218

**Authors:** Abdul M. Bhuiya, Siddhansh Agarwal, Carlos F. Ng, Melanie Ott, Sungmin Son, Daniel A. Fletcher

**Affiliations:** Department of Bioengineering, University of California, Berkeley, Berkeley, CA, United States; Biohub, San Francisco, CA 94158, USA; Gladstone Infectious Disease Institute, Gladstone Institutes, San Francisco, CA 94158, USA; Department of Medicine, University of California, San Francisco, San Francisco, CA 94158, USA; Department of Bio and Brain Engineering, KAIST, Daejeon, Republic of Korea; Division of Biological Systems and Engineering, Lawrence Berkeley National Laboratory, Berkeley, CA 94720, USA

## Abstract

CRISPR-based assays offer sensitive and specific nucleic acid detection for a broad range of diagnostic and surveillance applications. While standard solution-based assays are simple, they typically lack the ability to detect multiple targets individually without the need for more complex instrumentation and multi-step workflows. Surface-based strategies offer greater potential for multiplexing since the identity of a target can be associated with a location on a slide or a specific bead, as in the case of DNA arrays or Luminex bead assays. However, the design principles for optimizing surface-based CRISPR assays have yet to be developed. Here we present a multi-scale analytical framework integrating nanoscale enzyme kinetics, microscale diffusion constraints, and macroscale assay parameters that we use to develop a bead-based Cas13a assay where the bead becomes fluorescent when the target RNA for that bead is detected. The assay, which we call SurfCas, uses surface-bound guide RNAs and reporter RNAs to detect one or more soluble target RNAs in a one-pot assay. We show that SurfCas has a sensitivity approaching bulk reactions, and we demonstrate multiplexed detection of target RNAs in a complex biological matrix. The design principles we establish for surface-based CRISPR assays offer guidance on how to further improve SurfCas sensitivity and scale multiplexed detection of target RNAs in a single reaction.

## INTRODUCTION

Rapid, accurate, and scalable diagnostic methods are essential for timely pathogen detection, effective public health responses, and comprehensive disease surveillance. Conventional RNA detection techniques such as reverse transcription quantitative PCR (RT-qPCR) and next-generation sequencing offer high sensitivity but commonly require complex instrumentation, specialized training, and multiple steps that can introduce biases or errors^1–3^. Thus, there is an interest in developing RNA diagnostic platforms that combine amplification-free detection, operational simplicity, and multiplexing capabilities suitable for diverse clinical and field applications.

CRISPR-based diagnostics have recently emerged as a highly promising alternative to conventional diagnostic assays for RNA, offering programmable, sensitive, and sequence-specific nucleic acid detection without the target amplification requirements characteristic of PCR-based and sequencing methods^4^. These assays typically rely on reporter RNAs, which are single-stranded RNA molecules dual-labeled with a fluorophore and a quencher. Upon target recognition, Cas13 enzymes undergo a conformational change that triggers trans-cleavage of these reporters, releasing the fluorophores from their quenchers to generate a measurable fluorescent signal^5^. Multiplexing RNA detection typically requires orthogonal detection channels for each target to distinguish whether a target was detected, but an activated Cas13 enzyme will cut the same reporter RNA for any target RNA detected.

Using CRISPR-Cas enzymes in surface-based assays could enable multiplexed detection on beads or slides if the surfaces can effectively capture and concentrate the targets. The central challenge is designing the assay to maximize signal and minimize background, while accounting for diffusion, surface reactions, and sample volumes. CRISPR-based tests such as SHERLOCK already detect single RNA molecules with remarkable sensitivity, but practical multiplexing still stalls at only a handful of targets^6^. The limitation is not complexity; it is encoding capacity. CRISPR multiplexing normally requires orthogonal detection channels to distinguish which target was detected. Only a handful of Cas13 and Cas12 variants have sufficiently distinct nucleotide cleavage preferences for parallel detection, and fluorescent reporters face spectral overlap beyond 4–6 channels^7–11^. SHERLOCK variants achieve 4-target detection by exploiting orthogonal Cas enzymes but are constrained by enzymatic scarcity.

Overcoming these limitations involves fundamental trade-offs. The original CARMEN platform achieved impressive 169-virus detection by compartmentalizing reactions in thousands of nanoliter microfluidic droplets, but this approach requires splitting samples across many separate chambers^7^. Newer iterations such as mCARMEN for respiratory virus panels, bCARMEN for bacterial pathogens, and qCARMEN for quantitative mRNA profiling have adapted this technology for diverse clinical and research applications^12–14^. However, even these newer versions are constrained by a reliance on specialized microfluidic hardware and custom instrumentation, which limits their deployment in resource-constrained environments where such equipment is unavailable. In contrast, a fundamental advantage of our proposed system is sample conservation: instead of dividing clinical specimens across separate micro-compartments, all targets are detected simultaneously in a single, “one-pot” reaction. This one-pot configuration ensures that the entire sample volume is available to every sensor, maximizing the probability of detecting rare targets.

While bead-based platforms like Luminex routinely multiplex more than 50 targets using fluorescent barcoding, adapting them for CRISPR-based RNA detection assays is a relatively new endeavor^15^. Recent work has introduced bbCARMEN, which uses barcoded beads to enable parallelized detection of nine viral targets^16^. However, bbCARMEN still relies on pre-amplification steps—multiplexed RT-PCR or isothermal RPA—to achieve clinical sensitivity, which increases operational complexity and time-to-result. Additionally, the platform requires a complex workflow involving droplet formation and emulsification to isolate reactions, making it less suitable for rapid, one-step deployment. In contrast, our platform achieves multiplexed detection in a truly amplification-free, one-pot format, eliminating the need for both pre-processing steps and specialized microfluidic isolation.

Other existing bead-based CRISPR platforms have prioritized absolute sensitivity over multiplexing capacity, but they do so through significant trade-offs in workflow simplicity and scalability. For instance, the CRISPR-on-beads assay achieves high sensitivity for MRSA but requires isothermal pre-amplification and a complex, multi-step capture process, limiting its utility for broad diagnostic use^17^. Similar approaches have utilized magnetic beads to enhance bacterial and viral detection; however, these methods remain tied to multi-step, amplification-dependent workflows^18,19^. In a similar trade-off, a single-microbead Cas12a assay provides remarkable sensitivity without pre-amplification, yet it is fundamentally restricted to single-target detection and has complex preparation steps^20^. These systems lack a scalable barcoding architecture for “one-pot” multiplexing and often require specialized, high-resolution microscopy to resolve signals from individual beads, making them more suitable for niche single-molecule research than for high-throughput, multi-target diagnostics.

Our approach overcomes these constraints by using spectrally encoded fluorescent beads. Each target is assigned to a unique bead barcode. If the bead surface is functionalized with target-specific guide RNAs and a tethered reporter (fluorophore-quencher pair), every bead type can use the same Cas13 enzyme to locally cleave the reporter and generate a signal on the bead—orthogonality comes from bead identity, not molecular constraints. In this configuration, when Cas13 activates on bead A (containing guide RNAs for target A), local reporter concentrations are orders of magnitude higher than any reporters on distant bead types. Most cleavage occurs locally before the enzyme can diffuse and interact with other non-specific targets, such as reporters on neighboring beads or target RNA in the bulk solution. In addition, surface immobilization enables wash steps to remove unbound material, excess enzymes, and interfering substances. These capabilities are not possible in solution-based assays where everything remains mixed, and reactions continue until enzyme degradation.

However, surfaces introduce distinct challenges. Enzyme activity decreases upon tethering^16,17,20^. Mass transport limits reaction kinetics^21^. Surface chemistry creates bead-to-bead variability^22^. The vast parameter space makes empirical optimization prohibitively complex, perhaps explaining why CRISPR diagnostics have remained predominantly solution-based despite clear multiplexing advantages offered by bead-based platforms^23^. Here, we present SurfCas, a bead-based platform that uses magnetic, fluorescently barcoded streptavidin beads functionalized with biotinylated guide RNAs and fluorescent reporters for simultaneous detection of multiple RNA targets in a one-pot assay. Rather than empirical optimization, we designed SurfCas through comprehensive theoretical modeling that has resulted in a quantitative framework for surface-based CRISPR diagnostics. The modeling challenge required bridging four orders of magnitude: from millimeter-scale reaction geometries to micron-scale bead surfaces to nanometer-scale molecular interactions between tethered enzymes and substrates. We formulated coupled reaction-diffusion equations connecting bulk transport, surface binding kinetics, and constrained enzymatic activity—processes that cannot be decoupled experimentally.

This framework revealed why surface-based CRISPR assays have been challenging to optimize. A major obstacle has been background activity that compromises detection limits^23,24^. Our models identified two distinct sources with different kinetics: solution-phase Apo-Cas13 (lacking guide RNA), cleaving surface-tethered reporters, and surface-immobilized ribonucleoproteins cutting reporters even without targets present. Understanding these mechanisms allowed us to design targeted mitigation strategies for each. Central to our approach was developing an “accessibility factor” that quantifies how biotin-streptavidin tethering constrains Cas13-reporter interactions. This revealed a threshold linker length below which enzymatic activity is severely compromised, enabling rational design that ensures adequate accessibility. The models also predicted optimal parameters across other scales: bead sizes and numbers that balance capture efficiency against signal dilution, and optimal guide RNA-to-reporter ratios that maximize signal while controlling background. An interesting insight emerged about operational timing. The framework shows that incubation times >1 hr improve detection limits by 2-10 fold, but we selected 1-hour protocols for practical diagnostics while noting that extended times benefit surveillance applications where sample-to-answer time is less critical.

Experimental validation confirmed the predictive capability of our SurfCas modeling framework. Using only kinetic rates from independent time-lapse fluorescence experiments, the models accurately predicted optimal parameters without additional fitting. As a first demonstration of multiplexing, SurfCas achieves multiplexed detection of three targets with picomolar sensitivity in 1-hour assays—approaching solution-phase CRISPR performance for direct detection with a single guide, despite surface constraints. The framework extends beyond SurfCas to include amplified assays, establishing design principles for any surface-tethered enzymatic assay for different enzymes, chemistries, and detection modalities. This represents a methodological shift from empirical optimization to predictive design in surface-based diagnostics, combining CRISPR’s programmability with proven bead-based multiplexing to overcome fundamental constraints that have limited both technologies independently.

## MATERIALS AND METHODS

### Design of biotinylated guide RNAs and reporter RNAs

crRNA for SARS-CoV-2 sequences was designed as described in Fozouni et al. (2021) with a 20-nucleotide spacer and a 30-nucleotide stem. Reporters were designed as previously described in Fozouni et al. (2021) with a 5′ FITC and a 3′ Iowa Black quencher, connected through a 5U sequence. Several modifications to both crRNA and reporter were tested in order to tether them to the surface of beads. Biotin was added to either the 5′ end or the 3′ end of the crRNA; better Cas13 reaction activity was observed when the biotin was attached to the 3′ end (see Supplementary Figure S1). To optimize the assay, we evaluated various linker lengths for both the crRNAs and the reporters, including a no-spacer condition and spacers of approximately 12.6 nm and 50.4 nm, composed of single-stranded DNA and 18-atom hexa-ethyleneglycol units. For both crRNAs and reporters, a 20-molecule repeat of 18-atom hexa-ethyleneglycol spacers corresponding to an approximate linker length of 50.4 nm yielded the best Cas13 activity out of all the linkers tested (Supplementary Text Section 6 and Supplementary Table S1). The modified oligonucleotides were ordered from IDT. For crRNA containing a photocleavable molecule and an interfering DNA portion, an internal photocleavable spacer was added 5′ of the crRNA stem, which is cleaved upon exposure to 365 nm light dose as described in Ng et al. (2026)^25^, together with a 20T sequence added to the 5′ of the photocleavable spacer, which serves to block Cas13a activity as detailed in Son et al. (2026)^26^.

### LbuCas13a expression and purification

LbuCas13a enzyme was expressed and purified following previously established protocols^27^. Briefly, the enzyme was expressed in *E. coli* using a codon-optimized construct featuring an N-terminal His6-MBP-TEV fusion tag (Addgene Plasmid #83482). Following initial affinity purification and tag cleavage, the protein was further refined via size-exclusion chromatography using a S200 column (GE Healthcare). Purified fractions were collected in a HEPES-buffered storage solution containing 10% glycerol, flash-frozen, and stored at -80°C for subsequent assays.

### Preparation of color-coded streptavidin beads with surface-bound guide RNAs and reporter RNAs

COMPEL™ Uniform Magnetic Microspheres (Streptavidin, 8 µm; UMC0102) were employed for all standard experiments, while microspheres of 6 µm (UMC0101) and 3 µm (UMC0100) sizes were used specifically for bead-size comparison experiments (Fig. 3).

Cas13 and crRNA were first combined at equimolar concentrations (500 nM each) and incubated for 10 min at room temperature to form ribonucleoprotein (RNP) complexes. Prior to use, streptavidin-coated magnetic beads were vortexed and briefly sonicated in a water bath to ensure uniform dispersion. They were then diluted to achieve approximately 1,000 beads per experimental condition. The pre-assembled RNP complexes and biotinylated RNA reporters were subsequently added to the diluted bead suspension at defined molar ratios to achieve desired surface densities. To prevent unwanted reporter cleavage, Murine RNase inhibitor (NEB, M0314) was added during bead preparation at a concentration of 1 unit/µL. For a typical 1:1 surface ratio, 500 nM each of RNP and reporter were incubated with the beads.

After incubation, beads were separated magnetically and washed four times with 1 mL of 1X reaction buffer (20 mM HEPES-Na pH 7.2, 50 mM KCl, 5 mM MgCl2 and 5% glycerol). Prepared beads were either stored at 4 °C or kept at room temperature until use in SurfCas reactions.

### SurfCas reactions on beads with tethered guide RNAs and reporter RNAs

Since Cas13a can dissociate from the tethered guide RNA, 1 nM of Cas13 was added to the bead mixture and incubated for 10 minutes to equilibrate the amount of RNP on the bead surface. The reaction mixtures were then transferred to a glass-bottom 384 well plate (Corning, Cat# 4581). Synthetic target RNA or RNase-free water, along with 5X reaction buffer containing magnesium and 7.5% PEG-8000 (w/v), was then added to the bead mixture. The concentration of PEG-8000 was optimized to enhance reaction performance through macromolecular crowding, particularly for target concentrations near the limit of detection (see Supplementary Text Section 6 and Figure S3). Finally, the beads were mixed via pipetting to initiate the reaction, and the plate was sealed with an optically clear film (ThermoFisher Scientific, Cat# Ab1170) to prevent evaporation and allow microscopic imaging of the wells.

### Plasma sample preparation

Plasma samples were derived from single-donor human whole blood (Innovative Research) collected from a 44-year-old Caucasian female donor. To isolate the plasma, whole blood collected in Acid Citrate Dextrose (ACD) was centrifuged at 4,000 rpm for 10 minutes at 4°C. The resulting supernatant was aliquoted and stored at -80°C until use. For experiments, these plasma samples were diluted to 25% (v/v) in reaction buffer. Proteinase K (ThermoFisher Scientific, Cat# EO0491) was then added to the plasma at a final concentration of 50 µg/ml. The mixture was incubated at 55°C for 15 min, followed by a heat inactivation step at 95°C for 5 min. After cooling it down to room temperature, synthetic target RNA or RNase-free water (as a negative control) was spiked into the plasma-buffer mixture. The resulting mixture was then combined with the bead preparation for SurfCas analysis. Standard laboratory safety practices were followed during this process.

### Quantification of surface-tethered reporter molecules and RNP molecules

Quantum™ FITC-5 MESF beads were used to estimate the number of surface-bound reporter molecules, and Quantum™ Alexa Fluor® 647 MESF beads were used to estimate the number of surface-bound RNP molecules (see Supplementary Text Section 6 and Figure S2). These calibration beads, which contain a defined number of fluorophores, were used to generate a standard curve correlating the fluorescence signal obtained from the microscope with the number of fluorophores per bead under the specific imaging settings employed^28,29^.

### Imaging and analysis of beads in endpoint experiments

For endpoint imaging, a widefield fluorescence microscope (Nikon) equipped with a 20X/0.80 NA objective was used to image the beads. Multiple fields of view were acquired to capture the entire well. Beads located at the edge of the field of view and clusters comprising two or more beads were excluded from the measurements. A spot detection pipeline in NIS Elements was employed to segment the beads based on an intensity threshold and circularity, and circular regions of interest (ROIs) were drawn for each bead based on this segmentation. Images were corrected for illumination non-uniformity in the FITC channel by acquiring a corresponding “correction” image of the well plate without beads and using the shading correction pipeline in NIS Elements. The sum fluorescence intensity of each bead was measured from the ROIs. Background fluorescence from beads with tethered reporters and RNPs in the no-target control (NTC) was subtracted from the sum fluorescence measurements of beads with target RNA. When the NTC was included in a figure, background fluorescence from the surrounding solution was subtracted after illumination correction using ImageJ. All data were subsequently plotted in GraphPad Prism, and statistical comparisons were performed using two-tailed t-tests.

### Imaging and analysis of beads in time-lapse experiments

For time-lapse imaging to determine the kinetics of the Cas13 reaction, a confocal fluorescence microscope with a 40X objective was used to image the beads. Z-stack images spanning the entire diameter of the beads were captured for three fields of view per well at 30-second intervals. For experiments with photocleavable guide RNA, time-lapse images were captured in the FITC channel at 30-second intervals for 5 minutes, followed by a 30 second UV pulse at 365 nm (X-Cite 120Q Fluorescence Illuminator). This induces cleavage of the photocleavable portion of the guide RNA, releasing the interfering DNA segment and transitioning the enzyme from a suppressed to an activated state. After stimulation, time-lapse imaging was resumed for 1 hour at 30-second intervals. Images were corrected for photobleaching using the bleach correction plugin in ImageJ with an exponential fit method. In a separate condition, positive control beads with the same surface configuration as the experimental beads were treated with RNase A (ThermoFisher Scientific, Cat# EN0531) to cleave all reporters on the surface and then imaged over time with identical laser power, exposure time, and time intervals. These images were used to correct for photobleaching under the assumption that fluorescence on the positive control beads should remain constant over time. Tracking was performed in FIJI using the TrackMate plugin to monitor bead movement and record the corresponding signal for each bead. A spot detection pipeline in NIS Elements was again used to segment the beads based on an intensity threshold and circularity. Images were corrected for illumination non-uniformity in the FITC channel by acquiring a corresponding “correction” image of the well plate without beads and applying the shading correction pipeline in NIS Elements. The sum fluorescence intensity for each bead was measured from the ROIs, and background fluorescence from the surrounding solution was subtracted after illumination correction using ImageJ. All data were subsequently plotted in GraphPad Prism, and statistical comparisons were performed using two-tailed t-tests.

## RESULTS AND DISCUSSION

### SurfCas assay design

The SurfCas assay is based on the use of bead barcoding to identify specific target RNAs and fluorescent signal generation on the same bead surface to report if the target RNA has been detected (Fig. 1A). Initially, Cas13a enzymes are complexed with biotinylated guide RNAs at an equimolar ratio to form binary ribonucleoprotein (RNP) complexes in solution. These RNPs and biotinylated reporter RNAs (single-stranded RNAs with a fluorophore and a quencher on either end), each featuring long flexible linkers, are incubated with magnetic streptavidin-coated beads at defined molar ratios to functionalize the bead surface. Magnetic beads were employed to improve coupling efficiency while reducing reagent consumption and facilitating efficient washing through magnetic separation, as prior studies have shown^30,31^. The relative densities of these two components can be controlled by varying their input ratios during preparation. Upon addition of the Cas13a enzyme in solution, binary ribonucleoprotein (RNP) complexes form with the immobilized guide RNAs at the bead surface. When target RNA is present in the sample solution and encounters a bead surface, it hybridizes to the spacer regions of the bead-bound RNPs forming active ternary complexes. The activated Cas13a then initiates trans-cleavage of the nearby immobilized reporter RNAs, releasing the quencher molecules, resulting in a measurable increase in fluorescence at the bead surface (Fig. 1B).

**Fig. 1.**
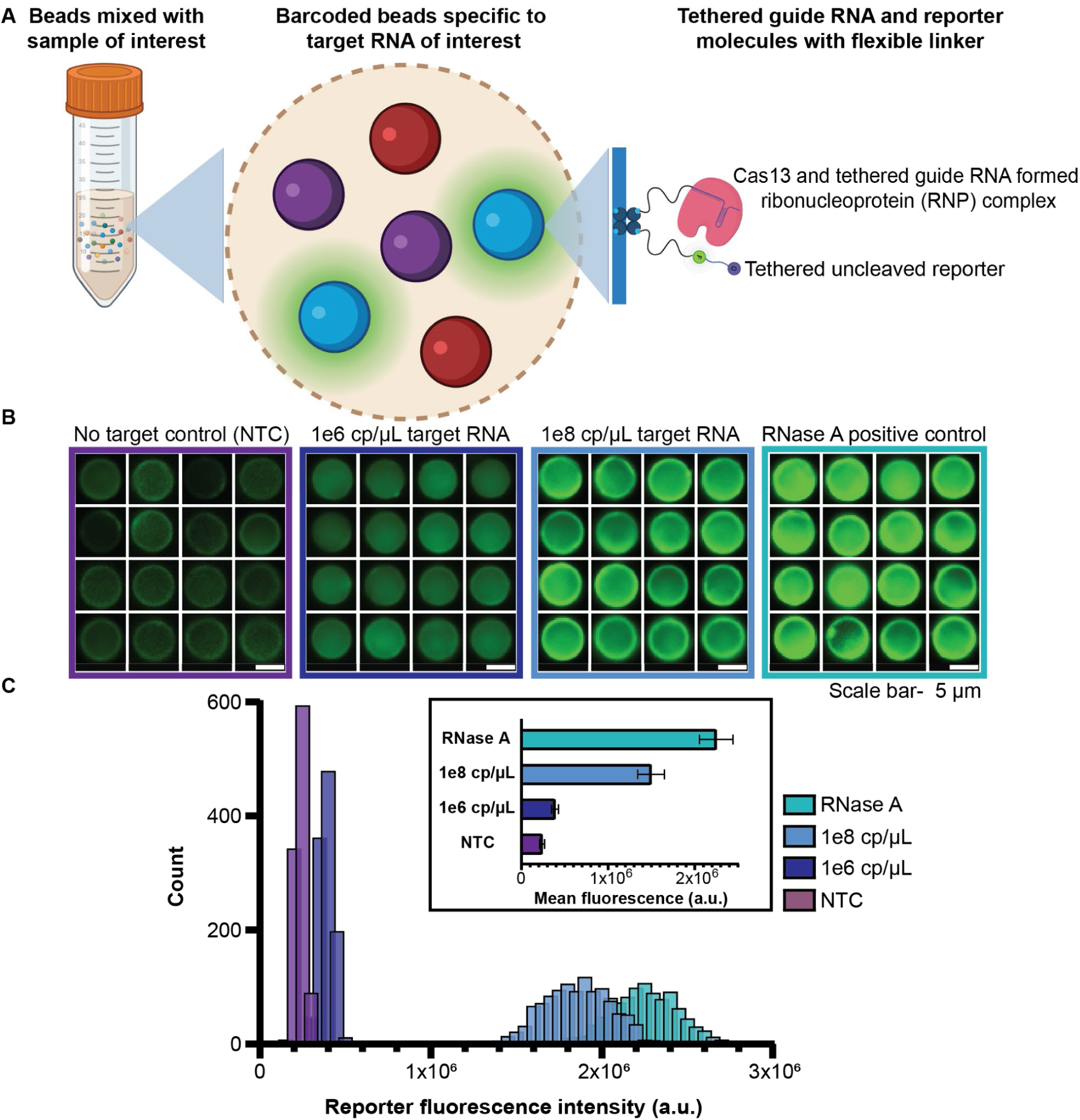
SurfCas reaction workflow, data visualization and analysis. (**A**) Schematic illustration of the SurfCas reaction workflow. (**B**) Epifluorescence microscopy images of beads representing negative control, positive control, and varying concentrations of spiked synthetic target RNA. (**C**) Fluorescence intensity distributions of beads corresponding to the experimental conditions shown in (B). Inset shows quantification of mean fluorescence intensities derived from the distributions in (C) across different target RNA conditions. Error bars represent the standard deviation of bead fluorescence intensities.

To quantify fluorescence of a population of beads detecting a specific RNA target using fluorescence imaging, regions of interest (ROIs) are drawn around each bead based on a mask from brightfield images of beads. Fluorescence intensity from each bead is calculated by summing pixel intensities within these ROIs, generating distributions of fluorescence intensities for different populations of beads (Fig. 1C). The mean fluorescence intensities from these distributions can be compared for different conditions and target RNA concentrations (Figure 1c inset), showing that the fluorescence signal increases with increasing target RNA concentration in solution.

### Multi-scale modeling framework

The exceptional molecular specificity of CRISPR-Cas systems offers tremendous potential for diagnostics. However, surface-based assays are difficult to optimize due to the challenge of coupled transport and reaction phenomena spanning six orders of magnitude in length scale. While molecular recognition between guide RNA and target occurs at nanometer scales, overall system performance depends critically on micron-scale diffusion around individual beads and millimeter-scale collective effects across entire bead populations in 10 µL-scale reaction volume. Understanding these multi-scale interactions is essential because optimization strategies that enhance performance at one scale may paradoxically degrade it at another. With this in mind, we developed a quantitative framework by recognizing that surface-based CRISPR diagnostics involve sequential reaction processes that must be coupled to diffusion. As shown in Fig. 2A, our approach partitions the sample volume *V_S_* containing N dispersed beads into identical spherical cells^21,32^, where each cell of radius *R* = (3*V_S_*/4π*N*)^1/3^ contains one bead at its center (Fig. 2A). This transforms the complex many-body problem into a tractable single-cell analysis while preserving collective depletion effects.

**Fig. 2.**
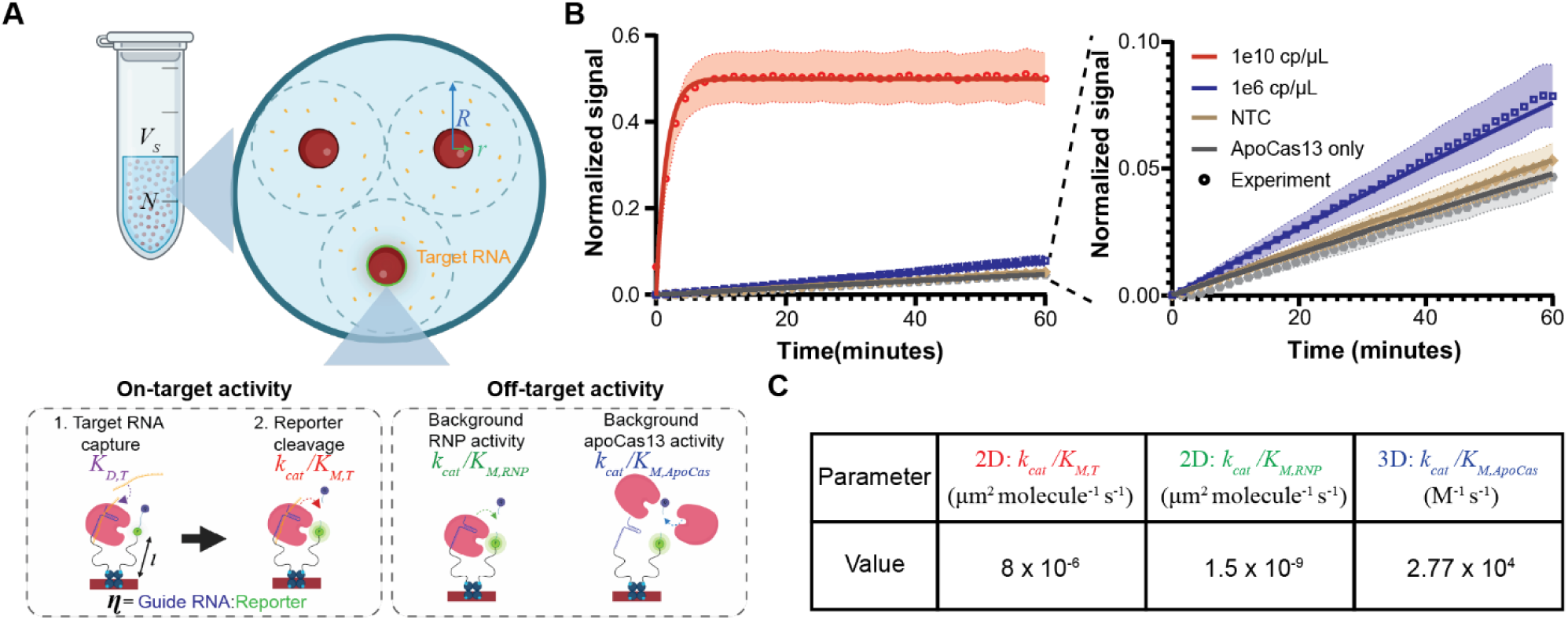
Analytical framework and characterization of SurfCas surface reaction kinetics. (**A**) Schematic depicting key parameters and reaction steps included in the SurfCas analytical framework. Parameters defined are: V_s_, reaction volume; N, Number of beads; r, radius of beads; R, radius of the spherical region around each bead defined by R = (3V_s_ /4πN)^1/3^; *K_D_*, dissociation constant; *K_CAT_*, catalytic rate constant for reporter cleavage; *K_M_*, Michaelis constant. (**B**) Mean bead fluorescence intensities during UV-synchronized time-lapse experiments using photocleavable guide RNAs, measured for target, apo-Cas13-only, and NTC conditions. Beads were imaged for 5 min before a 365 nm UV pulse to initiate guide uncaging, after which fluorescence was monitored for 1h. Traces are shown for spiked synthetic target RNA at 1e10 cp/µl and 1e6 cp/µl, along with apo-Cas13 only condition and an NTC condition. The apo-Cas13-only condition reflects solution-phase apo-Cas13 cleavage of surface-tethered reporters, whereas the NTC reflects total background in absence of target. Experimental data points are indicated by circles, and theoretical predictions from the analytical model are represented by solid lines. (**C**) Model parameters extracted from the time-lapse data in (B), distinguishing the 3D bulk apo-Cas13 background pathway from the 2D surface-tethered RNP pathway. For surface-tethered reactions, catalytic efficiencies are reported with respect to surface density rather than bulk concentration.

Within each cell, concentration fields of Cas enzyme [*E*] and target [*T*] are modeled by the diffusion equation in spherical coordinates: 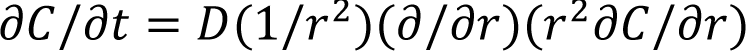, with boundary conditions at the bead surface (r = r₀) that couple nanoscale binding events to micron-scale concentration gradients:

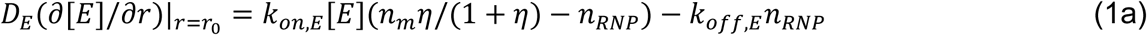

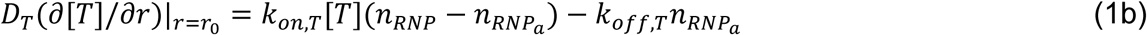

Here *n_m_* represents total surface density of guide-reporter pairs, η is the guide-to-reporter ratio on the bead surface, *n_RNP_* is the surface density of RNP complexes, *n*_*RNPa*_ represents activated RNP complexes capable of signal generation, and *D*_*E*_, *D*_*T*_ are respective diffusion coefficients. These boundary conditions capture how surface reactions consume nearby molecules, creating concentration gradients that drive further transport.

The system exhibits distinct timescales: Local diffusion around individual beads occurs on τ_*diff*_ = *r*_0_^2^/*D* ≈ 0.2 s (2 s) for target (enzyme), while molecular dissociation requires τ_*off*_ = 1/*k*_*off*_ ≈ 1 s. The concentration-dependent association time τ_*on*_ = 1/*k*_*on*_[*T*] varies dramatically from 17 minutes at 1 pM to nearly 12 days at 1 fM detection limits. Bulk homogenization across the bead ensemble requires τ_*bulk*_ = *r*^2^/*D* ≈ 3 mins (30 mins) for target (enzyme). This hierarchy means trace detection occurs far from binding equilibrium—local processes around beads equilibrate rapidly while bulk concentrations evolve slowly, allowing quasi-steady approximations for surface densities even during ongoing bulk depletion. Non-dimensionalizing with bead radius *r*₀, target dissociation constant *K*_*D,T*_ = *k*_*off,T*_/*k*_*on,T*_, and dissociation rate *k*_*off,T*_ reveals key dimensionless groups. The surface Damköhler number *D*_*a*_ = *n_m_r*_0_*k*_*on*_/*D* ≈ 10^5^ (10^6^) for target (enzyme) compares surface reaction rates to diffusive supply rates. Large *D*_*a*_ values indicate that reactions can outpace molecular transport, creating depletion zones around beads. The target depletion parameter λ_*T*_ = 4π*r*_0_^2^*Nn_m_*/(*V_S_K*_*D,T*_) quantifies when collective binding across all beads significantly reduces bulk concentrations. When λ_*T*_ ≪ 1, each bead operates independently in an effectively infinite reservoir. When λ_*T*_ ≫ 1, the beads collectively consume a significant fraction of available targets.

We use the quasi-steady approximation to gain analytical solutions for surface densities (see Supplementary Text Section 1 for complete derivation). The non–dimensional RNP surface density formation at steady-state follows 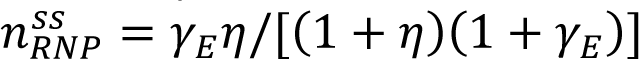, where Υ_*E*_ = [*E*]_0_/*K*_*D,E*_. This expression approaches the geometric limit η/(1 + η) when enzymes are abundant (Υ_*E*_ ≫ 1). Target capture becomes more complex due to potential bulk depletion. Mass balance requires 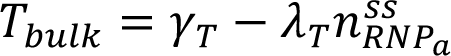, where Υ_*T*_ = [*T*]_0_/*K*_*D,T*_, yielding:

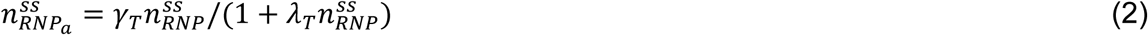

This expression reveals a key insight: the activated RNP surface density is proportional to target concentration Υ_*T*_, but with an efficiency factor determined by the target depletion parameter λ_*T*_. The denominator represents a reduction in capture efficiency due to target depletion, becoming significant when λ_*T*_ ≫ 1. Thus, surface localization provides a potential enrichment advantage when targets can be captured and reacted locally on beads without substantially reducing the bulk target pool.

### Predicted signal generation by the SurfCas assay

Signal generation by release of a fluorescent quencher from the reporter RNA must account for geometric constraints that fundamentally distinguish surface-tethered from solution-phase reactions. While solution-phase enzymes and substrates encounter each other through random diffusion, surface immobilization confines interactions to within combined tether reach, creating spatial constraints where productive encounters depend on molecular linker lengths and local density of active molecules. We capture this through an accessibility factor φ that models bidirectional geometric constraints—enzymes must find accessible substrates within their interaction radius defined by *l* = (*l*_*enzyme*_ +*l*_*reporter*_), where *l* is the combined linker length. As derived in Supplementary Text Section 2, this geometric constraint produces two distinct kinetic regimes. Near the limit of detection, activated complexes are sparse (*n*_*RNPa*_ ≪ *S*), and reporters must locate active enzymes through local search, creating quadratic signal dependence on activated RNP, independent of reporter density on the bead:

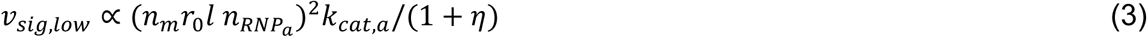

Signal amplification occurs through increased encounter probability as activated complexes accumulate. The quadratic appearance of combined linker length makes tether engineering a powerful sensitivity lever. When complexes become abundant, substrates easily find enzymes and the signal transitions to linear scaling with activated RNP density, approaching solution-phase behavior.

In our system, the background signal arises from two distinct sources with different kinetic expressions. Surface-bound RNP complexes exhibit weak target-independent cleavage with rate *v*_*bkg,RNP*_ ∝ *k*_*cat,RNP*_*n*_*RNP*_[*S*], where *k*_*cat,RNP*_ is the cleavage rate. In addition, apo-Cas13a (Cas13a without a guide RNA) in solution can cleave surface tethered reporter, creating background *v*_*bkg*,*Apo*_ ∝ *k*_*cat*,*Apo*_Υ_*E*_[*S*], where *k*_*cat*,*Apo*_ is the cutting rate. The dominant source determines optimal design parameters. Increasing the guide-to-reporter ratio η initially improves performance by enhancing signal generation through increased target capture, faster than background accumulation. However, higher η eventually worsens performance by accumulating more background signal, creating an optimum near η ≈ 3 in the RNP-dominated case and around η ≈ 1-2 in the apoCas13a-dominated situation. These optima are concentration-regime dependent and should be interpreted for the intended operating range rather than as universal settings.

To experimentally determine these kinetic parameters and validate our theoretical predictions, we performed time-lapse fluorescence measurements using photocleavable guide RNAs (Fig. 2B). The photocleavable guide RNAs contain a short photocleavable DNA oligo extended from the 5′ end of the crRNA, which is cleaved upon exposure to 365 nm UV light, allowing precise temporal control over reaction initiation^25^. This approach enabled accurate measurement of reaction kinetics by synchronizing the start of Cas13a activity across all beads. We measured fluorescence time-courses at high (1×10¹⁰ copies/µL) and low (1×10⁶ copies/µL) target RNA concentrations, along with an apo-Cas13-only control to isolate the bulk apo-Cas13 contribution and a no-target control capturing the net background of the full surface assay in the absence of target. By fitting our analytical model to these experimental time-courses, we extracted key parameters including catalytic rate constants, background activity rates for both RNP and apo-Cas13, and accessibility coefficients (Fig. 2C). These experimentally determined parameters were then used without further fitting to generate all subsequent model predictions for assay optimization.

Detection limits emerge from the fundamental requirement that target-specific signal must exceed background noise with statistical confidence^23,24^. In an ideal Poissonian system, the standard deviation equals the square root of the mean. However, real experimental systems exhibit additional noise sources beyond Poissonian statistics—including bead variability, optical noise, and systematic drift—that must be characterized from actual measurements. We employ a mean + 3SD criterion, where 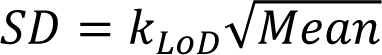 based on experimentally determined *k*_*LoD*_ (see Supplementary Text Section 2). The detection limit analysis reveals how the multi-scale physics of our system constrains sensitivity. In non-depleted conditions (λ_*T*_ ≪ 1), the limit of detection scales favorably with surface density as *n_m_*^−3/4^, improves with larger bead size as *r*_0_^−1/2^ and longer tethers as *l*^−1^, and benefits from higher catalytic rate as 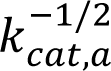. The background signal depends critically on which mechanism dominates—RNP-mediated or apo-Cas13a-mediated cleavage—determining optimal guide-to-reporter ratios (complete expressions in Supplementary Text Sections 2–3). However, in depleted conditions (λ_*T*_ ≫ 1), the limit of detection worsens as *Nr*_0_^3/2^*n*_m_^1/4^/*V_S_* ; although increasing surface area can improve capture in the non-depleted regime, adding more beads or increasing their size beyond the depletion threshold can paradoxically worsen performance by enhancing background faster than improving capture efficiency.

Our framework reveals that nanoscale recognition, micron-scale transport, and millimeter-scale collective effects create emergent behaviors not obvious from molecular properties alone. The quadratic scaling at low concentrations provides unique sensitivity advantages through surface geometry, but only with proper tether optimization. System performance depends critically on identifying the operating regime—depleted versus non-depleted—and the dominant background source. This multi-scale understanding enables rational optimization by revealing the fundamental physical mechanisms and trade-offs governing bead-based CRISPR diagnostic performance.

### Experimental validation of multi-scale model predictions

To test the predictions of our theoretical framework, we systematically varied key parameters and measured NTC-subtracted signal across multiple target concentrations (Fig. 3 and Supplementary Text Sections 2 and 6). Our bead number and radius experiments (Fig. 3A and Fig. 3B) directly demonstrate the depletion transition predicted by our framework. At high target concentration (10¹⁰ copies/μL), the mean fluorescence signal increased predictably with total surface area *Nr*_0_^2^, consistent with non-depleted operation where depletion is negligible (λ_*T*_ ≪ 1). However, at lower concentrations (10⁶ copies/μL), the mean fluorescence signal initially increased with total surface area, indicating a beneficial surface-capture regime, but then peaked at an intermediate value and declined rapidly after crossing the predicted depletion threshold. This behavior occurs when λ_*T*_ exceeds unity, where collective binding across beads begins depleting bulk target concentrations faster than individual beads can capture them. The threshold condition *Nr*_0_^2^ < *V_S_K*_*D,T*_(1 + η)(1 + Υ_*E*_)/(4π*n_m_* Υ_*E*_η) establishes a quantitative upper limit on optimal surface area. Beyond this limit, detection worsens as target molecules distribute across more binding sites while the background signal continues to increase proportionally with surface area. This finding explains why simply adding more capture surface can fail at low concentrations—the very regime where sensitivity matters most.

**Fig. 3.**
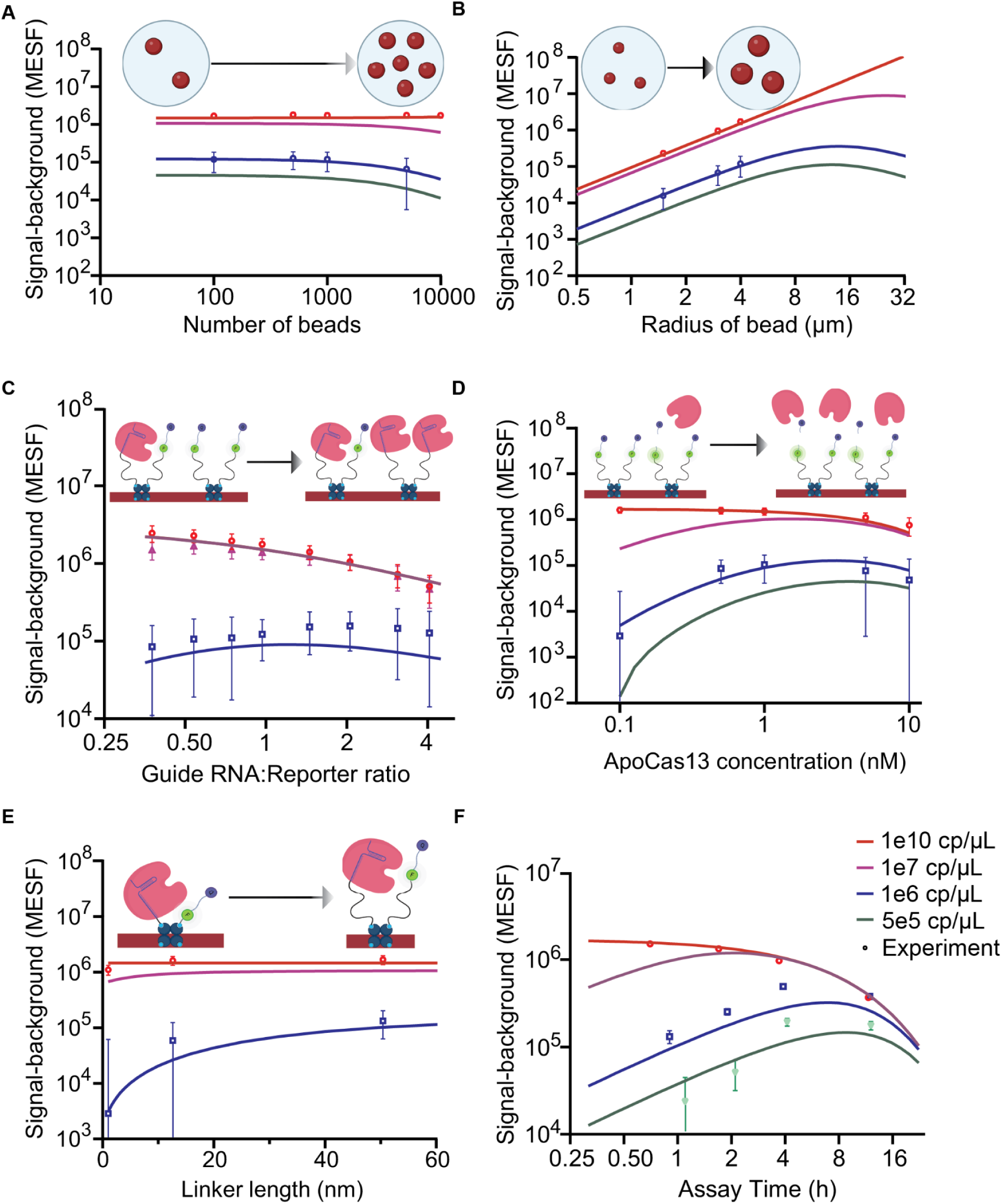
SurfCas model predictions and experimental validation of key parameters. (**A-B**) Model predictions and experimental validation for macroscale parameters: (**A**) bead number and (**B**) bead size. At low total surface area, increasing surface area improves signal by enhancing local target capture on beads. Beyond the depletion threshold, further increases in surface area worsen performance because bulk targets are depleted across too many binding sites while background continues to accumulate. (**C–E**) Model predictions and experimental validation for nanoscale parameters: (**C**) guide RNA-to-reporter ratio, (**D**) apoCas13 concentration, and (**E**) linker length. Error bars represent standard deviation. Error bars are truncated where values would fall below zero in log-log scale. (**F**) Model predictions and experimental validation for operational parameter: Assay time. Experimental data points are indicated by circles, and theoretical predictions from the analytical model are represented by solid lines.

The guide-to-reporter ratio experiments (Fig. 3C) reveal surface density-dependent optimal values that further confirm our analysis. At high target concentrations (10¹⁰ copies/μL), lower η values maximize signal by providing more reporters to convert abundant activated complexes into detectable output. As target concentration decreases (10⁶ copies/μL), the optimum shifts higher because target capture becomes limiting instead of signal conversion. Our experimental system, dominated by apo-Cas13a background, showed optimal η ≈ 1-2 for lower concentrations, matching theoretical curves derived from numerical solutions of our complete model (Fig. 3D and detailed in Supplementary Text Section 2).

Molecular tether length experiments (Fig. 3E) support our accessibility model and the predicted quadratic dependence on combined linker length. Signal enhancement with increasing tether length was most pronounced at low target concentrations, with an observed saturation at approximately 50 nm in length. Assay time optimization (Fig. 3F) confirmed our prediction that optimal measurement duration depends on target concentration through the balance between signal accumulation and background growth. At high target concentrations (10¹⁰ copies/μL), maximum signal-to-background ratio occurred at the shortest time tested (30 minutes), after which it decreased as reactions approached completion while background continued accumulating. At lower concentrations (10⁷ to 5×10⁵ copies/μL), optimal performance occurred between 2 and 12 hours, where neither signal nor background had saturated and the balance favored continued accumulation. Thus, we find that optimal measurement time scales inversely with target abundance.

These experiments demonstrate that our multi-scale modeling framework successfully captures the complex interplay between nanoscale recognition, micron-scale transport, and millimeter-scale collective effects, making it a useful tool for surface-based Cas13 assay optimization. The quantitative agreement between theory and experiment across multiple parameter variations validates several key insights: (i) depletion parameters determine operating regimes, (ii) surface geometry creates unique scaling advantages, (iii) background source identification governs optimal ratios, and (iv) time-resolved measurements are essential for sensitivity optimization.

### SurfCas assay direct detection limits

With optimized key parameters for SurfCas, we then systematically characterized the sensitivity of SurfCas by determining the limit of detection (LOD) when directly detecting target RNAs without amplification. We considered different tethered reaction configurations and used a guide RNA designed to detect synthetic SARS-CoV-2 target RNA in a 1-hour reaction (Fig. 4). When either the reporter RNA alone or the guide RNA alone are tethered to the bead, with the other reagents in solution, the LODs are approximately 5×10⁵ copies/µL (Figs. 4A and 4B), compared to a bulk reaction sensitivity of 1×10⁵ copies/µL for all reagents in solution. With both the guide RNA and reporter RNA tethered to beads, we achieved an LOD of approximately 1×10⁶ copies/µL (Fig. 4C). As expected, this shows that tethering any one reagent to the bead reduces sensitivity, while tethering two reagents has an additive reduction in sensitivity.

**Fig. 4.**
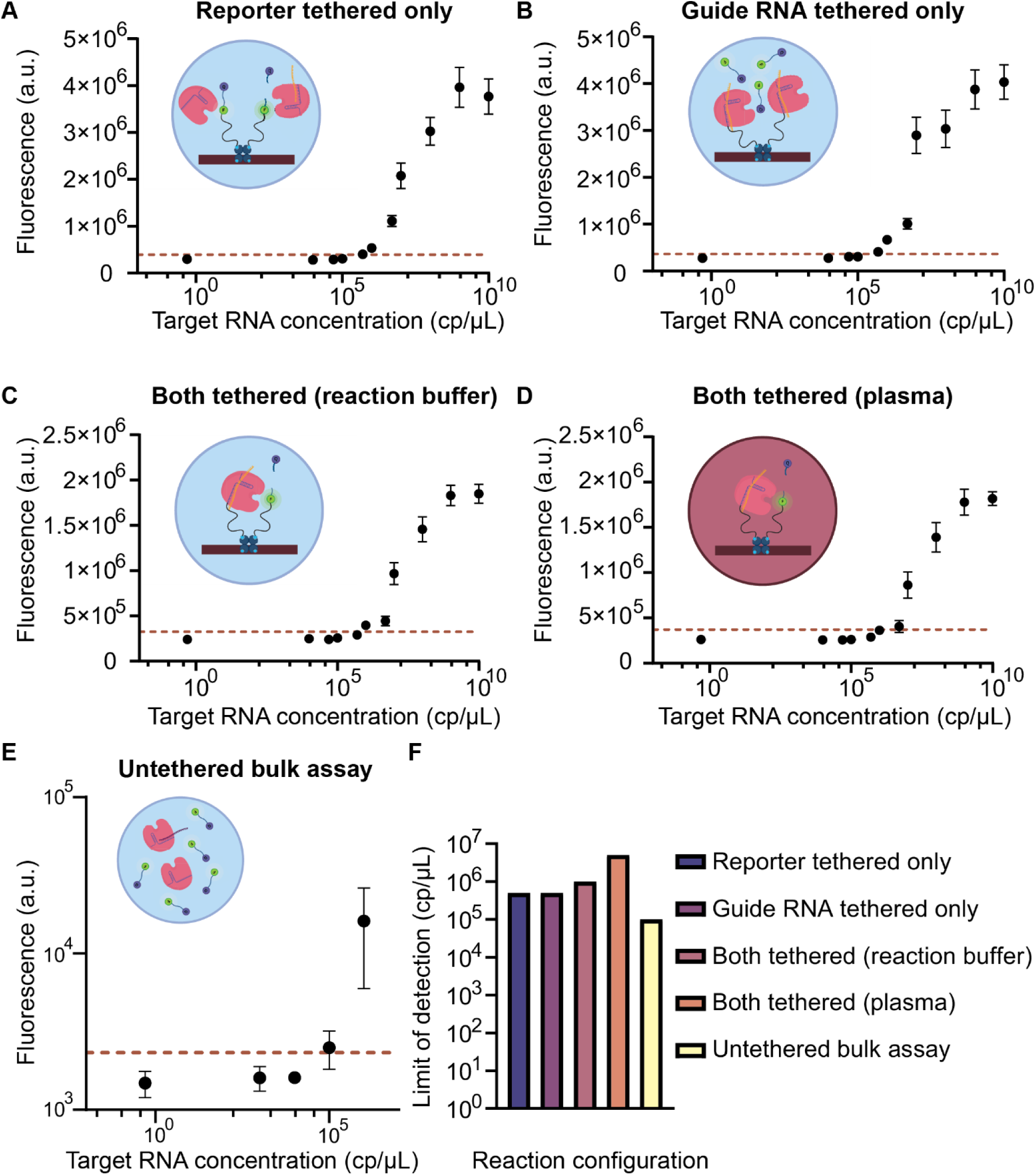
SurfCas experimental limits of detection. (**A-E**) Limit of detection (LOD) of SurfCas using optimized conditions with synthetic SARS-CoV-2 target RNA in reaction buffer with (**A**) guide RNA tethered only, (**B**) reporter tethered only, (**C**) both guide RNA and reporter tethered in reaction buffer; solid line represents theoretical prediction from the analytical model, (**D**) both guide RNA and reporter tethered in plasma and (**E**) bulk assay where both guide RNA and reporter are not tethered to any surface. Error bars indicate the standard deviation of three replicates. Red dashed lines indicate the limit of detection (LOD), defined as the mean of the replicates of no-template control (NTC) condition plus three times the standard deviation of replicates (mean NTC + 3 × SD). (**F**) Summary plot comparing the LOD for all the different reaction configurations

To further evaluate assay performance in clinically relevant conditions, we determined the LOD for synthetic SARS-CoV-2 RNA spiked into plasma samples. Under these conditions, the dual tethered assay achieved an LOD of 5×10⁶ copies/µL for a 1-hour reaction (Fig. 4D). This sensitivity is reduced but comparable to results obtained in buffer, demonstrating the robustness of the SurfCas assay in complex biological matrices. The dual-tethered bead-based configuration approached the sensitivity of the bulk assay, differing by only an order of magnitude, demonstrating that our platform retains reasonable sensitivity in a surface-bound format (Fig. 4F). To better understand the parameters governing these limits and identify pathways for further optimization, we turned to an analytical modeling framework.

We leveraged our analytical framework to predict the limit of detection (LOD) of our current system, incorporating both background sources—RNP and apo-Cas13a activities—as well as an idealized scenario without apo-Cas13a background activity (Fig. 5A and 5B). Our analysis highlights that optimal LOD depends strongly on both RNP background activity (x-axis) and apo-Cas13a background activity (y-axis). Critically, our predictions indicate that removing the apo-Cas13a background reduces the LOD by roughly an order of magnitude. The alignment between predicted curves and experimental data across varying conditions confirms our model’s predictive power, enabling quantitative insights into the sensitivity-speed trade-offs essential for rational assay optimization tailored to specific diagnostic applications.

**Fig. 5.**
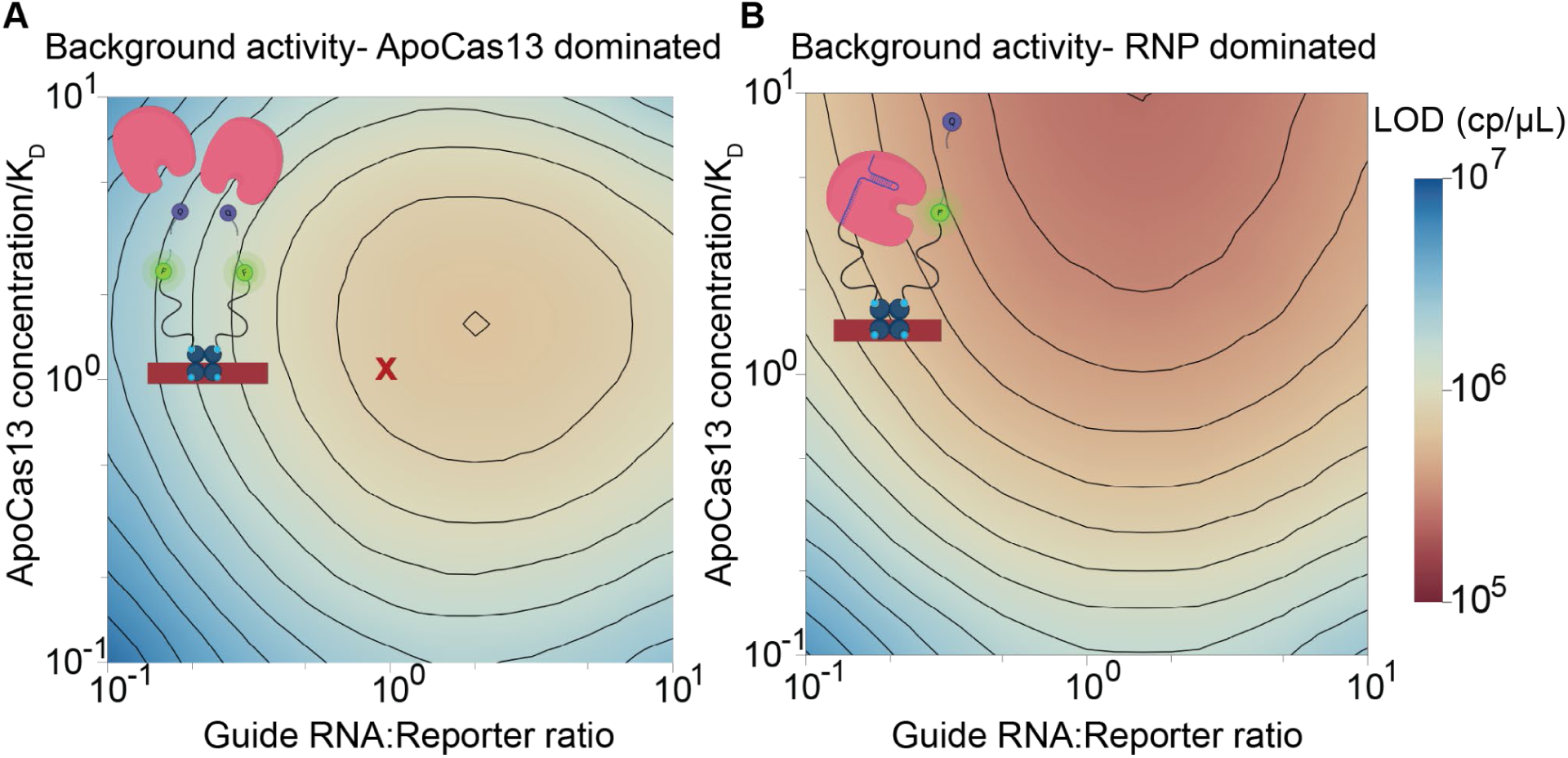
SurfCas predicted limit of detection. (**A-B**) Model predictions for SurfCas limit of detection (LOD) under two different background activity conditions: (**A**) with both RNP and apoCas13 background activity, and (**B**) with apoCas13 background activity only. The red X in (A) denotes the LOD for our current SurfCas system.

### Multiplexed detection with SurfCas in one-pot assay

With the sensitivity of the assay well characterized, we next demonstrated the assay’s capacity for simultaneous multi-target detection. This was achieved in a one-pot format by utilizing distinct fluorescently barcoded beads mixed in a single reaction volume to detect multiple RNA targets. Specifically, three distinct bead populations were functionalized with low concentrations of fluorescent biotin, using emission wavelengths that avoid spectral overlap with the reporter RNA channel. This strategy enabled unique color-coding of distinct bead populations without negatively impacting reaction kinetics (Fig. 6A (i)). Each fluorescently labeled bead population was tethered with specific guide RNA-reporter pairs targeting three distinct regions of the SARS-CoV-2 genome: two within the N gene and one within the Orf1ab gene. We spiked 1×10^7^ copies/µL of each synthetic target RNA into three reaction mixtures, each comprising all three bead populations. Fluorescence microscopy confirmed selective activation only in bead populations matched to their respective synthetic RNA targets (Fig. 5A (ii)). For example, “blue-coded” beads functionalized with crRNA 4 targeting the SARS-CoV-2 N gene exhibited fluorescence exclusively when synthetic RNA containing the corresponding N gene sequence was introduced, confirming the specificity of our multiplexing strategy.

**Fig. 6.**
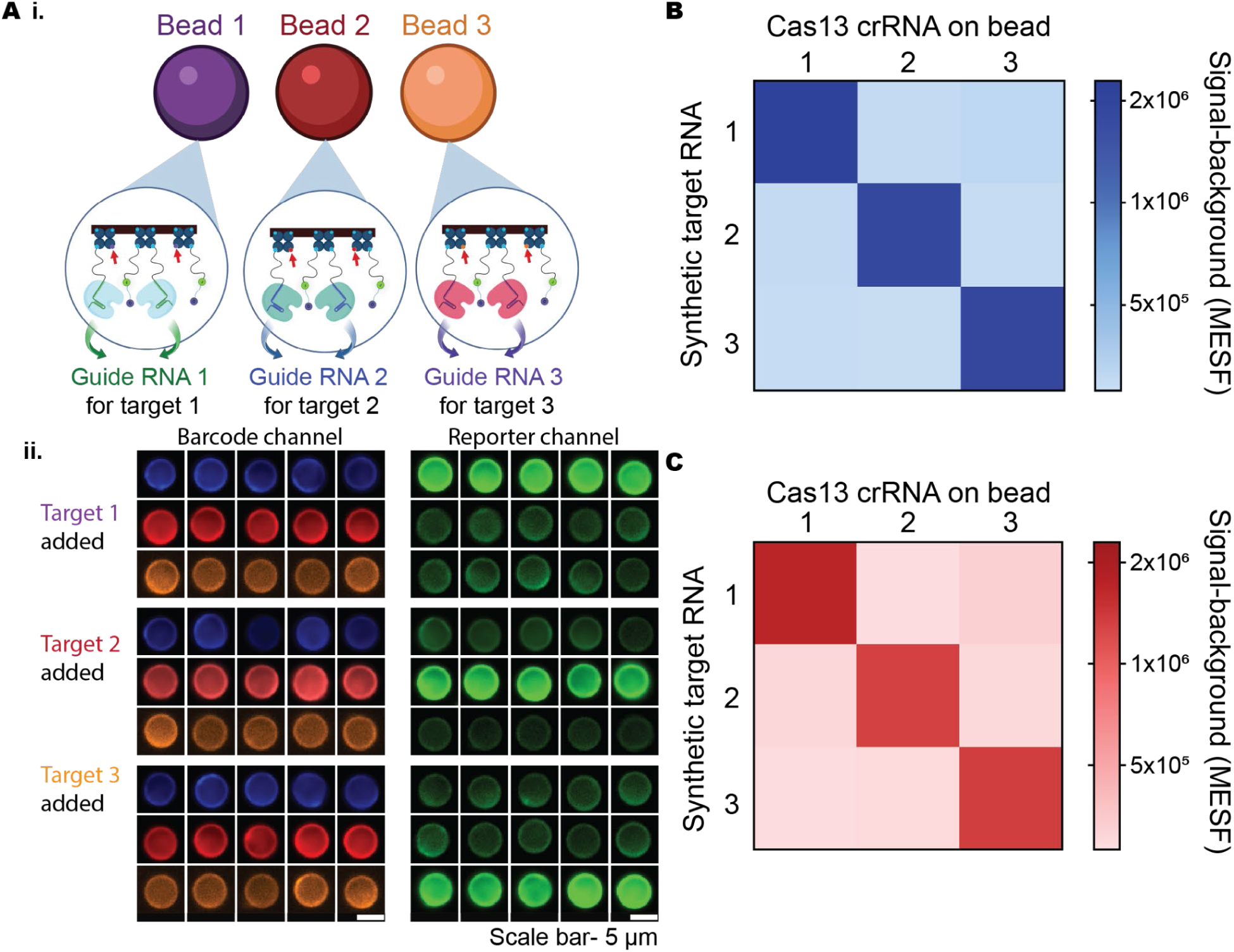
SurfCas multiplexing performance in complex biological matrices. (**A**) Multiplexing demonstration: (**i**) schematic illustration of the multiplexing strategy; (**ii**) epifluorescence microscopy images of fluorescently barcoded beads, each tethered with unique guide RNAs and challenged with their respective target RNAs. Images show fluorescence in barcode and reporter channels. (**B**) Quantification of mean fluorescence reporter intensities from beads in (**A** (**ii**)), demonstrating high specificity and minimal cross-reactivity in reaction buffer. (**C**) Quantification of mean fluorescence reporter intensities from multiplexed bead assays in plasma, confirming specificity and minimal off-target activity under biologically relevant conditions.

Quantitative validation of multiplex performance was carried out using synthetic RNA targets representing the distinct SARS-CoV-2 genomic regions. The resulting fluorescence intensity heatmap demonstrates high specificity (Fig. 6B), with high signals corresponding only to matched guide RNA-target RNA pairs. Multiplexed detection of target RNA was also carried out in plasma, further confirming assay specificity within biologically complex samples (Fig. 6C). These findings highlight our assay’s robust discrimination capability in multiplexed formats.

## CONCLUSIONS

In this study, we have demonstrated a novel bead-based Cas13 diagnostic platform, which we call SurfCas, that integrates a comprehensive multi-scale modeling framework with a one-pot assay format compatible with highly scalable multiplexing. Our approach addresses several critical challenges in translating CRISPR-based detection to field-ready applications. By preassembling Cas13 reaction components on barcoded streptavidin magnetic beads, our system minimizes sample handling and obviates the need for complex instrumentation. The one-pot format simplifies the assay workflow, reducing potential user error and significantly shortening turnaround time. This streamlined approach is particularly beneficial in resource-limited settings. Our work bridges a critical gap between molecular-scale engineering and system-level performance in diagnostic platforms by establishing quantitative physical constraints that govern detection sensitivity. The remarkable agreement between theoretical predictions and experimental results over several orders of magnitude in target concentration (10^5^-10^10^ copies/μL) supports our multi-scale approach while revealing previously unrecognized limitations in conventional diagnostic design. Perhaps most surprisingly, we found that maximizing binding capacity, which might initially appear to be a good choice, can undermine sensitivity for trace analytes. This counterintuitive effect occurs when systems cross the depletion threshold 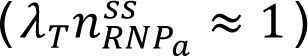, where target molecules become distributed across too many binding sites. For our experimental conditions, this establishes a specific upper bound (*Nr*_0_^2^ < 2.5 × 10^−8^ m²) on optimal surface area—explaining inconsistencies in reported sensitivities across various biosensing platforms and providing a quantitative foundation for rational design decisions.

Surface confinement fundamentally alters reaction kinetics through geometric constraints that are not present in solution-phase systems. Our accessibility model captures these effects, accurately predicting both the quadratic dependence on linker length and signal saturation observed at ∼50nm length. This mathematical framework explains why molecular motion constraints on surfaces create unique scaling relationships that determine detection limits, particularly at low target concentrations where accessibility becomes rate-limiting. The optimal guide-to-reporter ratio (η=1-2) and temporal relationship (*t*_*opt*_ ≈ (1/*k_sig_*) ⋅ ln[(*k_sig_* + *k_bkg_*)/*k_bkg_*]) further illustrate how molecular design choices must account for specific physical trade-offs. These quantitative relationships provide precise guidelines for optimizing parameters based on application requirements, whether prioritizing ultimate sensitivity or rapid detection.

The analytical framework we have developed extends naturally to diverse contexts. For systems with lateral diffusion, such as lipid-tethered molecules or cell membrane receptors, our accessibility factor adapts by replacing fixed tether lengths with appropriate diffusion length scales (√Dt). The depletion parameter can incorporate flow rates and channel geometries for microfluidic implementations, while the background component can be modified for systems with different dominant noise mechanisms.

Several promising innovations emerge from our theoretical insights. Enzyme immobilization strategies using oriented attachment (e.g., via SpyTag/SpyCatcher, maleimide chemistry, or click chemistry) could dramatically reduce apo-Cas13a background while maintaining catalytic activity—potentially improving detection limits by 1-2 orders of magnitude through altered background mechanisms^33–35^. Similarly, DNA origami scaffolds could enhance molecular accessibility beyond what random-coil linkers achieve by creating precisely positioned 3D arrangements of reaction components. Furthermore, incorporating multiple distinct guide RNAs on each bead, targeting different regions of the pathogen genome, can enhance sensitivity, as previously demonstrated by Fozouni et al. and further confirmed for the SurfCas configuration in this study (see Supplementary Text Section 6 and Supplementary Figure S4)^27^. Beyond these optimizations, the effective sensitivity of SurfCas could be further augmented by incorporating pre-amplification of nucleic acid (e.g. RPA or LAMP) prior to bead-based readout. This strategy, like the approach employed by the CARMEN platforms, would significantly lower the limit of detection and allow detection of ultra-low abundance targets. However, such an implementation would necessitate a trade-off, introducing additional liquid-handling steps and operational complexity that may impact the platform’s suitability for simple field deployment.

Another major advantage of a bead-based platform is its inherent capacity for multiplex detection. By assigning distinct guide RNAs to uniquely barcoded beads, we have successfully demonstrated the simultaneous detection of three RNA targets. This strategy not only improves diagnostic throughput but also offers the potential to expand the assay to cover broader panels of pathogens or biomarkers, an essential feature for comprehensive diagnostic and surveillance applications. Future work will focus on refining reaction conditions, expanding the target panel, and integrating portable imaging systems for real-time readout. Additionally, extensive field validation in diverse sample matrices such as clinical specimens and environmental samples will be essential to demonstrate the utility of this platform. In summary, SurfCas represents an important step forward in the development of rapid, robust, and multiplexed nucleic acid tests. By combining a user-friendly one-pot assay with a rigorous theoretical framework, we offer a promising blueprint for next-generation diagnostic tools capable of meeting the diverse demands of modern public health challenges.

## ASSOCIATED CONTENT

**Supporting Information:** The Supporting Information is available free of charge and attached as a separate pdf.

## AUTHOR INFORMATION

### Author contributions

A.M.B.: Conceptualization, Methodology, Data curation, Formal analysis, Investigation, Experimental validation, Modeling framework, Visualization, Writing – original draft; S.A.: Conceptualization, Modeling framework, Methodology, Data curation, Formal analysis, Visualization, Writing – original draft; C.F.N.: Investigation, Experimental validation, Writing – review & editing; S.S.: Conceptualization, Writing – review & editing; M.O.: Funding acquisition, Resources, Supervision, Writing – review & editing; D.A.F.: Conceptualization, Funding acquisition, Resources, Supervision, Project administration, Writing – review & editing.

## ACKNOWLEDGEMENTS

We thank all members of the Fletcher and Ott laboratories for critical discussions and feedback on this project. We acknowledge BioRender for creation of schematic for all of the figures in the publication.

