## Supplemental Information for "Bead-based RNA Detection with Cas13a: Multi-scale Design and Validation"

**This PDF File Includes**
Supplementary Text
Supplementary Figures S1 to S4
Supplementary Table S1
Supplementary References

### Supplementary Text

#### 1. Theoretical Framework: Model Formulation and Dimensionless Analysis

Our bead-based CRISPR diagnostic system presents a complex multi-scale problem that we approach through reaction-diffusion modeling. The system consists of spherical beads with radius $r_0$ dispersed throughout a sample volume $V_s$. Each bead carries surface-immobilized guide RNAs and quenched fluorescent reporters in a ratio $\eta$:1. Rather than attempting to solve the full many-body problem, we assume each bead sits at the center of a spherical region with radius $R = (3V_s/4\pi N)^{1/3}$. This approach divides the sample into $N$ identical regions, each containing exactly one bead, which works well for experimentally realistic bead dispersions. Within each cell (spanning from $r_0$ to $R$), we track two mobile species: Cas13a with no guide RNA or target RNA (Apo-Cas) in solution molecules $[E]$ and target RNA molecules $[T]$. Both species diffuse according to Fick's second law in spherical coordinates:

$$\frac{\partial C}{\partial t} = D\frac{1}{r^2}\frac{\partial}{\partial r}\left(r^2\frac{\partial C}{\partial r}\right) \quad \text{for } C \in \{[E],[T]\}. \tag{S1}$$

The interesting chemistry happens at the bead surface ($r = r_0$). Here, Apo-Cas enzymes bind to surface-anchored guide RNAs, forming ribonucleoprotein (RNP) complexes with surface density $n_{\text{RNP}}$. These RNP complexes then capture target molecules in solution to create activated complexes with density $n_{\text{RNPa}}$. We couple the bulk diffusion to these surface reactions through flux-kinetic boundary conditions:

$$D_E\frac{\partial[E]}{\partial r}\bigg|_{r=r_0} = k_{\text{on},E}[E](r_0)\left(\frac{n_m\eta}{1+\eta} - n_{\text{RNP}}\right) - k_{\text{off},E}n_{\text{RNP}} \tag{S2a}$$

$$D_T\frac{\partial[T]}{\partial r}\bigg|_{r=r_0} = k_{\text{on},T}[T](r_0)(n_{\text{RNP}} - n_{\text{RNPa}}) - k_{\text{off},T}n_{\text{RNPa}} \tag{S2b}$$

In these equations, $n_m$ represents the total surface density of binding sites, and the factor $\eta/(1+\eta)$ gives the fraction occupied by guides. At the outer boundary of each cell, we impose zero-flux condition, $\frac{\partial C}{\partial r}\Big|_{r=R} = 0$ for $C \in \{[E],[T]\}$ since the system has symmetry around each bead.

**Dimensional Analysis and Characteristic Scales.** The system involves parameters spanning multiple orders of magnitude, making dimensional analysis essential for understanding the underlying physics. We choose four characteristic scales that capture the essential features: bead radius $r_0$ for length, target dissociation time $1/k_{\text{off},T}$ for time, target dissociation constant $K_{D,T} = k_{\text{off},T}/k_{\text{on},T}$ for concentration, and maximum bead surface density $n_m$ for surface coverage. This non-dimensionalization reveals several key parameter groups. The binding affinities are characterized by $\rho = K_{D,E}/K_{D,T}$, which compares how strongly enzymes versus targets bind to their respective surface sites. Here, we assume $K_{D,E} = K_{D,T} = K_D$ so that $\rho = 1$. The initial concentrations become $\gamma_E = [E]_0/K_D$ and $\gamma_T = [T]_0/K_D$. Denoting non-dimensional variables with a hat ($\hat{}$), the resulting non-dimensional equations are as follows:

$$1 < \hat{r} < \alpha: \quad \frac{\partial \hat{E}}{\partial \hat{t}} = \tau_E \nabla^2 \hat{E}, \quad \frac{\partial \hat{T}}{\partial \hat{t}} = \tau_T \nabla^2 \hat{T} \tag{S3a}$$

$$\hat{r} = 1: \quad \frac{1}{Da_E} \frac{\partial \hat{E}}{\partial \hat{r}}\bigg|_{\hat{r}=1} = \frac{\partial \hat{n}_{\mathrm{RNP}}}{\partial \hat{t}} = \hat{E}\left(\frac{\eta}{1+\eta} - \hat{n}_{\mathrm{RNP}}\right) - \hat{n}_{\mathrm{RNP}},$$

$$\frac{1}{Da_T} \frac{\partial \hat{T}}{\partial \hat{r}}\bigg|_{\hat{r}=1} = \frac{\partial \hat{n}_{\mathrm{RNP}_a}}{\partial \hat{t}} = \hat{T}(\hat{n}_{\mathrm{RNP}} - \hat{n}_{\mathrm{RNP}_a}) - \hat{n}_{\mathrm{RNP}_a} \tag{S3b}$$

$$\hat{r} = \alpha: \quad \frac{\partial \hat{E}}{\partial \hat{r}} = 0, \quad \frac{\partial \hat{T}}{\partial \hat{r}} = 0 \tag{S3c}$$

$$\hat{t} = 0: \quad \hat{E} = \gamma_E, \quad \hat{n}_{\mathrm{RNP}} = 0, \quad \hat{T} = \gamma_T, \quad \hat{n}_{\mathrm{RNP}_a} = 0 \tag{S3d}$$

Here, $\alpha = R/r_0$ is the nondimensional outer radius, $\tau_i = D_i/(k_{\mathrm{off},T} r_0^2)$ is the non-dimensional diffusion time. The competition between reaction and diffusion appears through Damköhler numbers $\mathrm{Da}_i = n_m r_0 k_{\mathrm{on},i}/D_i$. These represent the ratio of maximum reaction rate to maximum diffusion rate around each bead. For realistic diagnostic parameters, we find remarkably high values that fundamentally change how the system behaves, as discussed below.

**Transport-Reaction Balance and Timescale Analysis.** Let's examine what these Damköhler numbers mean in practice. Using typical values for surface density ($n_m = 10^4$ molecules/$\mu$m$^2$), bead radius ($r_0 = 4\mu$m), and association rates ($k_{\mathrm{on}} = 10^9$ M$^{-1}$s$^{-1}$), we calculate $\mathrm{Da}_E \approx 6.64 \times 10^6$ for enzymes and $\mathrm{Da}_T \approx 6.64 \times 10^5$ for targets. The difference arises because larger Cas enzymes (around 150 kDa) diffuse more slowly ($D_E \approx 10^{-11}$ m$^2$/s) than smaller nucleic acid targets ($D_T \approx 10^{-10}$ m$^2$/s). These extremely high Damköhler numbers tell us that both species are severely diffusion-limited. Molecules get consumed by surface reactions almost the instant they reach the bead surface, creating steep concentration gradients in the surrounding fluid. The local concentration right at the bead surface drops far below the bulk value, effectively reducing reaction rates from their intrinsic kinetic values to much slower diffusion-limited rates.

The system also exhibits a fascinating hierarchy of timescales. Local diffusion around individual beads happens quickly: enzymes take about 1.6 seconds to diffuse across a bead radius, while targets need only 0.16 seconds. Molecular dissociation occurs on similar timescales, typically around 1 second for both species assuming $k_{\mathrm{off}} = 1$ s$^{-1}$. But bulk transport across entire Wigner-Seitz cells takes much longer. For a typical experimental setup with 1000 beads in 20 $\mu$L, each cell has radius $R \approx 170\mu$m. Homogenizing enzyme concentrations across this distance requires about 48 minutes, while targets need roughly 5 minutes. This creates an interesting dynamic where each bead exists in a quasi-steady local environment that slowly evolves as the bulk concentrations change. The most dramatic timescale effects appear at low target concentrations. The association time $t_{\mathrm{react,on},T} = 1/(k_{\mathrm{on},T}[T])$ becomes painfully long when target concentrations drop. At 1 pM target concentration, this time is about 17 minutes. At 1 fM, it stretches to nearly 12 days. These extended association times mean that near detection limits, the system operates in a transient binding regime rather than reaching equilibrium during typical measurement periods.

**Surface Binding Dynamics and Steady-State Behavior.** Given the complex timescale hierarchy, we need approximations to make analytical progress. The high Damköhler numbers justify using a quasi-steady state (QSS) approximation for surface complexes, assuming they equilibrate rapidly compared to bulk concentration changes. We also assume negligible enzyme depletion, which holds when enzyme concentrations are high, enzyme affinity is weak, or total surface area is small relative to sample volume. The QSS assumption sets the time derivatives to zero:

$$\hat{E}(\hat{r}=1)\left(\frac{\eta}{1+\eta}-\hat{n}^{\rm ss}_{\rm RNP}\right)-\hat{n}^{\rm ss}_{\rm RNP}=0, \quad \hat{T}(\hat{r}=1)(\hat{n}^{\rm ss}_{\rm RNP}-\hat{n}^{\rm ss}_{\rm RNPa})-\hat{n}^{\rm ss}_{\rm RNPa}=0 \tag{S4}$$

We solve for $\hat{n}^{\rm ss}_{\rm RNP}$ and $\hat{n}^{\rm ss}_{\rm RNPa}$ to obtain:

$$\hat{n}^{\rm ss}_{\rm RNP}=\frac{\hat{E}(\hat{r}=1)\frac{\eta}{1+\eta}}{1+\hat{E}(\hat{r}=1)}, \quad \hat{n}^{\rm ss}_{\rm RNPa}=\frac{\hat{T}(\hat{r}=1)\hat{n}^{\rm ss}_{\rm RNP}}{1+\hat{T}(\hat{r}=1)} \tag{S5}$$

The concentration at the bead surface is related to the bulk concentration via mass balance equations: $V[E]_0 = V[E] + 4\pi r_0^2 N n_{\rm RNP}$ and $V[T]_0 = V[T] + 4\pi r_0^2 N n_{{\rm RNP}_a}$, which in non-dimensional form read:

$$\hat{E}(\hat{r}=1)\approx\hat{E}_{\rm bulk}=\gamma_E-\lambda_E\hat{n}^{\rm ss}_{\rm RNP}, \quad \hat{T}(\hat{r}=1)\approx\hat{T}_{\rm bulk}=\gamma_T-\lambda_T\hat{n}^{\rm ss}_{\rm RNPa}. \tag{S6}$$

We obtain the depletion parameter $\lambda_i = 4\pi r_0^2 N n_m/(V_s K_{D,i})$, which measures the ratio of total binding capacity across all beads to the number of target (or enzyme) molecules available at the dissociation constant concentration. This parameter ultimately determines whether the system operates in a favorable regime where molecular optimizations matter, or in a depleted regime where adding more binding capacity actually hurts performance. Substituting Equation S6 into Equation S5 and rearranging yields quadratic equations. When enzyme depletion is negligible ($\lambda_E \to 0$), this simplifies to:

$$\hat{n}^{\rm ss}_{\rm RNP}=\frac{\gamma_E\eta}{(1+\eta)(1+\gamma_E)} \tag{S7}$$

This expression shows how RNP formation depends on both enzyme concentration and surface stoichiometry. As enzyme concentration increases, the RNP density approaches a maximum value of $\eta/(1+\eta)$, which represents the fraction of surface sites available for RNP formation.

Target binding presents a more complex situation because target depletion can be significant even at low concentrations. The QSS solution for the activated complex formation takes the form of a quadratic equation:

$$\hat{n}^{\rm ss}_{\rm RNPa}=\frac{(1+\gamma_T+\lambda_T\hat{n}^{\rm ss}_{\rm RNP})-\sqrt{(1+\gamma_T+\lambda_T\hat{n}^{\rm ss}_{\rm RNP})^2-4\lambda_T\gamma_T\hat{n}^{\rm ss}_{\rm RNP}}}{2\lambda_T} \tag{S8}$$

For low target concentrations ($\gamma_T \ll 1$), which matter most near detection limits, this simplifies to:

$$\hat{n}_{\text{RNPa}}^{\text{ss}} \approx \frac{\gamma_T \hat{n}_{\text{RNP}}^{\text{ss}}}{1 + \lambda_T \hat{n}_{\text{RNP}}^{\text{ss}}} \tag{S9}$$

This expression captures a fundamental design trade-off. The activated complex density scales linearly with target concentration, but the efficiency factor $1/(1 + \lambda_T \hat{n}_{\text{RNP}}^{\text{ss}})$ creates two distinct operating regimes. When $\lambda_T \hat{n}_{\text{RNP}}^{\text{ss}} \ll 1$, we're in the non-depleted regime where molecular-scale optimizations can dramatically improve sensitivity. But when $\lambda_T \hat{n}_{\text{RNP}}^{\text{ss}} \gg 1$, we enter the depleted regime where adding more binding capacity actually makes things worse.

**Signal Generation Mechanisms and Reporter Kinetics.** The ultimate goal is generating detectable fluorescent signal through reporter cleavage by activated RNP complexes. A critical distinction between solution-phase and surface-based enzymatic reactions lies in the geometric constraints of tethered molecules. For systems where both enzymes (activated RNP) and substrates (fluorescent reporters) are immobilized on a surface with no lateral diffusion, traditional reaction kinetics must be modified to account for fixed-position spatial constraints. Unlike freely diffusing molecules that can encounter each other through random motion, surface-tethered molecules have fixed positions: reactions can only occur between pairs that happen to be positioned within each other's reach. To model this, we develop an accessibility model that accounts for the bidirectional nature of the geometric constraint. While there can be many functional forms, we choose the following for the accessibility factor:

$$\phi = \frac{1}{\frac{(1+\eta)}{\eta}\frac{\hat{n}_{\text{RNPa}}^{\text{ss}}}{\hat{S}}\frac{1}{\beta_e} + \frac{(1+\eta)\hat{S}}{\hat{n}_{\text{RNPa}}^{\text{ss}}}\frac{1}{\beta_s} + 1} \tag{S10}$$

where $\beta_e = c_e \cdot \pi l^2 \cdot n_m$ and $\beta_s = c_s \cdot \pi l^2 \cdot n_m$ represent the maximum number of potential interaction partners within reach of an enzyme or substrate, respectively. Here $l$ is the combined tether length $l = l_{\text{enzyme}} + l_{\text{reporter}}$. The coefficients $c_e$ and $c_s$ account for the different geometric and orientation constraints in each direction and are obtained from experiments.

This accessibility formulation behaves like Michaelis-Menten kinetics but with important differences. In the enzyme-limited regime ($\hat{n}_{\text{RNPa}}^{\text{ss}} \ll \hat{S}$), the accessibility factor scales as $\hat{n}_{\text{RNPa}}^{\text{ss}}/\hat{S}$, making the overall signal quadratic in activated complex density. In the substrate-limited regime ($\hat{S} \ll \hat{n}_{\text{RNPa}}^{\text{ss}}$), it scales as $\hat{S}/\hat{n}_{\text{RNPa}}^{\text{ss}}$, giving linear signal dependence. When neither species is limiting, the accessibility approaches a constant determined by the surface geometry. The combined linker length ($l_{\text{enzyme}} + l_{\text{reporter}}$) appears squared in both $\beta_e$ and $\beta_s$, making linker engineering a powerful tool for sensitivity enhancement. Longer, more flexible linkers increase the interaction probability between activated complexes and reporters, though practical limits exist due to increased cross-talk and non-specific interactions.

After non-dimensionalizing using our characteristic scales, the dimensionless signal rate density becomes:

$$V_{\text{sig}} = -d\hat{S}/dt = \hat{k}_{\text{cat},T}\phi \hat{n}_{\text{RNPa}}^{\text{ss}}\hat{S} \tag{S11}$$

where $\hat{k}_{\text{cat},T} = n_m k_{\text{off},T} k_{\text{cat},T}$ is the non-dimensional surface cleavage rate constant and $\phi$ is our accessibility factor from Equation S10. The total background rate density combines contributions from unactivated RNP and Apo-Cas13 sources:

$$V_{\text{bkg}} = \left(\hat{k}_{\text{cat,RNP}}\hat{n}^{\text{ss}}_{\text{RNP}} + \hat{k}_{\text{cat,Apo}}\hat{E}_{\text{apo}}\right)\hat{S} \tag{S12}$$

Using the explicit form of $\hat{n}^{\text{ss}}_{\text{RNP}}$ from Equation S7 and with the initial reporter concentration $\hat{S}(0) = 1/(1+\eta)$, the total initial background rate can be expressed as:

$$B'(\eta) = \left(\hat{k}_{\text{cat,RNP}}\left[\frac{\gamma_E \eta}{(1+\eta)(1+\gamma_E)}\right] + \hat{k}_{\text{cat,Apo}}\gamma_E\right)\frac{1}{1+\eta} \tag{S13}$$

This expression reveals how background rates depend on enzyme concentration ($\gamma_E$) and surface stoichiometry ($\eta$). The relative contribution of each background term has significant implications for optimal system design, particularly for determining the optimal guide-to-reporter ratio $\eta$.

#### 2. Analytical Expressions for Diagnostic Performance

Translating our mechanistic understanding into practical performance metrics requires connecting surface binding dynamics to measurable fluorescence signals. The experimentally relevant quantity is the differential signal between target-containing samples and no-target controls: $\Delta F_{\text{bead}} = F_{\text{total,bead}}(\gamma_T) - F_{\text{total,bead}}(0) \approx (4\pi r_0^2) n_m V_{\text{sig}} k_{\text{off},T}\Delta t$.

**Target-Specific Signal Generation.** Using the initial rate approximation, the target-specific signal accumulated over measurement time $\Delta t$, in the low target limit, becomes:

$$\Delta F_{\text{bead}}(\gamma_T) = \frac{4\pi^2 c_s r_0^2 n_m^2 l^2 k_{\text{cat},T}}{(1+\eta)}\left(\frac{\gamma_T \hat{n}^{\text{ss}}_{\text{RNP}}}{1+\lambda_T \hat{n}^{\text{ss}}_{\text{RNP}}}\right)^2 k_{\text{off},T}\Delta t \tag{S14}$$

This expression encapsulates the entire pathway from target binding to signal generation. The quadratic dependence on activated complex density arises from our accessibility model and distinguishes surface-based diagnostics from solution-phase assays. The signal scales with bead surface area ($4\pi r_0^2$), total surface density squared ($n_m^2$), and combined linker length squared, highlighting the importance of engineering these physical parameters. For high target concentrations where accessibility constraints become less important, the signal transitions to linear scaling with activated complex density:

$$\Delta F_{\text{high}}(\gamma_T) \approx \frac{4\pi r_0^2 n_m k_{\text{cat},T}}{(1+\eta)}\hat{n}^{\text{ss}}_{\text{RNPa}} k_{\text{off},T}\Delta t \tag{S15}$$

**Limit of Detection Analysis.** The limit of detection represents the minimum target concentration that produces a signal distinguishable from background noise with specified statistical confidence. We use the standard criterion: $\Delta F_{\text{bead}}(\gamma_{T,\text{LoD}}) = k_{\text{LoD}}\sqrt{F_{\text{bkg,bead}}}$, where $k_{\text{LoD}} = 3$ for purely Poisson-limited detection. However, experimental noise sources like bead heterogeneity, instrumental fluctuations, and background variations increase this value to $k_{\text{LoD}} \approx 100$ in our system. Background signal arises from unactivated RNP complexes that retain some catalytic activity and free Apo-Cas enzymes that can cleave reporters non-specifically. We collect these contributions into $B'(\eta)$, which depends on the guide-to-reporter ratio through different mechanisms. Using our LoD criterion and solving for $\gamma_{T,\mathrm{LoD}}$ yields:

$$\gamma_{T,\mathrm{LoD}} = \frac{[T]_{\mathrm{LoD}}}{K_D} = \left(\frac{1+\lambda_T \hat{n}_{\mathrm{RNP}}^{\mathrm{ss}}}{\hat{n}_{\mathrm{RNP}}^{\mathrm{ss}}}\right)\sqrt{\frac{(1+\eta)}{k_{\mathrm{cat},T}\pi c_s l^2 n_m}}\left(\frac{k_{\mathrm{LoD}}^2 B'(\eta)}{4\pi r_0^2 n_m \Delta t k_{\mathrm{off},T}}\right)^{1/4} \tag{S16}$$

In the non-depleted regime where $\lambda_T \hat{n}_{\mathrm{RNP}}^{\mathrm{ss}} \ll 1$, molecular-scale parameters directly control sensitivity:

$$\gamma_{T,\mathrm{LoD}} = \frac{[T]_{\mathrm{LoD}}}{K_D} = \frac{(1+\eta)(1+\gamma_E)}{\gamma_E \eta}\sqrt{\frac{(1+\eta)}{k_{\mathrm{cat},T}\pi c_s l^2 n_m}}\left(\frac{k_{\mathrm{LoD}}^2 B'(\eta)}{4\pi r_0^2 n_m \Delta t k_{\mathrm{off},T}}\right)^{1/4} \tag{S17}$$

The LoD scales directly with the target-RNP dissociation constant $K_D$, with weaker binding leading to proportionally higher detection limits. Surface density appears as $n_m^{-3/4}$, so increasing surface coverage dramatically improves sensitivity. Combined linker length enters as $l^{-1}$, making longer linkers beneficial. Catalytic efficiency affects sensitivity as $(k_{\mathrm{cat},T})^{-1/2}$, while bead radius scales as $r_0^{-1/2}$. The fourth-root dependence on background makes the system relatively robust against background variations. In the depleted regime where $\lambda_T \hat{n}_{\mathrm{RNP}}^{\mathrm{ss}} \gg 1$, the story changes completely:

$$\gamma_{T,\mathrm{LoD}} = \frac{4\pi r_0^2 N n_m}{V_s K_{D,T}}\sqrt{\frac{(1+\eta)}{k_{\mathrm{cat},T}\pi c_s l^2 n_m}}\left(\frac{k_{\mathrm{LoD}}^2 B'(\eta)}{4\pi r_0^2 n_m \Delta t k_{\mathrm{off},T}}\right)^{1/4} \tag{S18}$$

Here, sensitivity worsens proportionally to $N \cdot r_0^{3/2} \cdot n_m^{1/4}$. Adding more beads, using larger beads, or increasing surface density all hurt performance because they enhance target depletion faster than they improve signal generation. This counterintuitive result explains why more isn't always better in diagnostic design. This depletion-driven trade-off is highlighted in the main-text experimental validation (Fig. 3A-B), where increasing available surface area improves signal at high target concentration but can reduce performance near the detection limit.

Following the same derivation approach, the LoD with perfect accessibility (or at high target concentrations) becomes:

$$\gamma_{T,\mathrm{LoD,perfect}} = \left(\frac{1+\lambda_T \hat{n}_{\mathrm{RNP}}^{\mathrm{ss}}}{\hat{n}_{\mathrm{RNP}}^{\mathrm{ss}}}\right)\frac{(1+\eta)}{k_{\mathrm{cat},T}} \cdot \sqrt{\frac{k_{\mathrm{LoD}}^2 B'(\eta)}{4\pi r_0^2 n_m \Delta t k_{\mathrm{off},T}}} \tag{S19}$$

Comparing equations S16 and S19 reveals two key insights. First, with perfect accessibility, LoD would scale with $\sqrt{B'(\eta)}$ rather than $(B'(\eta))^{1/4}$, making the system more sensitive to background reduction strategies. Second, linker engineering would no longer improve sensitivity—a direct consequence of removing spatial constraints. For the perfect accessibility case, the LoD in the non-depleted regime becomes:

$$\gamma_{T,\mathrm{LoD}} = \frac{(1+\gamma_E)}{\gamma_E \eta}\frac{(1+\eta)^2}{k_{\mathrm{cat},T}} \cdot \sqrt{\frac{k_{\mathrm{LoD}}^2 B'(\eta)}{4\pi r_0^2 n_m \Delta t k_{\mathrm{off},T}}}. \tag{S20}$$

In the depleted regime, it reads,

$$\gamma_{T,\mathrm{LoD}} = \frac{4\pi r_0^2 N n_m}{V_s K_{D,T}}\frac{(1+\eta)}{k_{\mathrm{cat},T}} \cdot \sqrt{\frac{k_{\mathrm{LoD}}^2 B'(\eta)}{4\pi r_0^2 n_m \Delta t k_{\mathrm{off},T}}} \tag{S21}$$

Even with perfect accessibility, the fundamental limitation from target depletion persists—the LoD still scales with $N \cdot r_0 \cdot n_m^{1/2}$. The key to achieving ultra-high sensitivity therefore lies in controlling depletion through careful selection of bead size, number, and sample volume, thereby creating conditions where molecular-scale optimizations can realize their full potential.

**Parameter Optimization for Enhanced Sensitivity.** Our analytical framework enables systematic optimization across the complex parameter space. The transition between operating regimes occurs when $\lambda_T \hat{n}_{\mathrm{RNP}}^{\mathrm{ss}} \approx 1$, establishing a fundamental design constraint:

$$N r_0^2 < \frac{V_s K_{D,T}}{4\pi n_m} \cdot \frac{(1+\eta)(1+\gamma_E)}{\gamma_E \eta} \tag{S22}$$

This simple relationship provides design guidance for balancing signal generation against depletion effects across various operating conditions. For high-sensitivity applications targeting trace target analytes, this relationship typically favors fewer, smaller beads to avoid depletion, while higher target-concentration assays can benefit from greater surface area.

Surface stoichiometry, represented by the guide-to-reporter ratio $\eta$, affects both signal generation and background noise through multiple mechanisms. The signal depends on $\eta$ through RNP formation ($\hat{n}_{\mathrm{RNP}}^{\mathrm{ss}} \propto \frac{\eta}{(1+\eta)}$), reporter density ($\propto \frac{1}{1+\eta}$), and accessibility effects. Simultaneously, different background sources exhibit distinct $\eta$-dependencies. The optimal guide-to-reporter ratio ($\eta$) depends on which background mechanism dominates in the system. For diagnostics where background primarily comes from unactivated RNP complexes, the $\eta$-dependence follows $\gamma_{T,\mathrm{LoD}}(\eta) \propto (1+\eta)/\eta^{3/4}$. Minimization through differentiation reveals that $\eta = 3$ provides optimal sensitivity in this scenario. When non-specific activity from free Cas enzymes (Apo-Cas) dominates the background, the relationship becomes $\gamma_{T,\mathrm{LoD}}(\eta) \propto (1+\eta)^{5/4}/\eta$, yielding an optimal value of $\eta = 4$. This slightly higher optimal ratio occurs because additional guide RNA sites improve signal generation without proportionally increasing Apo-Cas background. For systems where constant background sources predominate, we predict an optimal value of $\eta = 2$. These optimal values shift when we consider a perfect accessibility scenario ($\phi = 1$) where surface-tethered molecules can interact without geometric constraints. Under perfect accessibility conditions, the $\eta$-dependence becomes $\gamma_{T,\mathrm{LoD}}(\eta) \propto (1+\eta)/\eta^{1/2}$ for RNP-dominated background, $\gamma_{T,\mathrm{LoD}}(\eta) \propto (1+\eta)^{3/2}/\eta$ for Apo-Cas dominated background, and $\gamma_{T,\mathrm{LoD}}(\eta) \propto (1+\eta)^2/\eta$ for constant background. These relationships yield optimal $\eta$ values of 1 for both RNP and constant background scenarios, which aligns with intuitive expectations from a purely signal generation perspective. For Apo-Cas dominated systems, $\eta = 2$ remains the optimal value. In practical implementations, the accessibility factor satisfies $0 < \phi < 1$ depending on surface density, linker properties, and molecular orientation. Consequently, real-world diagnostic designs should typically maintain $\eta$ values between 1–2 for most applications, with higher values (approaching 3) beneficial primarily when Apo-Cas background specifically dominates and accessibility constraints are significant. This optimization represents an important trade-off between maximizing target capture efficiency and maintaining sufficient reporter density for effective signal generation. Importantly, these optima are operating-regime specific and should not be interpreted as universal across all target concentrations. Conditions that are favorable near the LoD can differ from those favored at high target abundance.

The measurement time presents another critical optimization parameter. The signal accumulation profile approximately follows:

$$\Delta F(t) \approx 4\pi r_0^2 n_m \frac{1}{1+\eta} \left( e^{-k_{\text{bkg}} t} - e^{-(k_{\text{sig}}+k_{\text{bkg}})t} \right), \tag{S23}$$

where $k_{\text{sig}} \propto \hat{n}_{\text{RNPa}}^{\text{ss}}$. Differentiating with respect to time and setting to 0, we find that the optimal measurement time is:

$$t_{\text{opt}} \approx \frac{1}{k_{\text{sig}}} \cdot \ln \left( \frac{k_{\text{sig}} + k_{\text{bkg}}}{k_{\text{bkg}}} \right). \tag{S24}$$

Since $k_{\text{sig}} \propto \gamma_T$ (to first order), the optimal time decreases approximately inversely with target concentration. This explains why higher target concentrations can be detected more rapidly, while low concentrations require extended measurement periods to achieve optimal sensitivity.

#### 3. Parameter Quantification and Integrated Optimization Strategy

To demonstrate practical application, consider a representative diagnostic system with 4 $\mu$m radius beads, 1000 beads dispersed in 20 $\mu$L sample volume, binding constants of 1 nM, surface density of $10^4$ molecules/$\mu$m$^2$, and enzyme concentration of 1 nM. This configuration gives $\gamma_E = 1$ and $\lambda_T \approx 0.167$. For $\eta = 1$, the steady-state RNP density becomes $\hat{n}_{\text{RNP}}^{\text{ss}} = 0.25$, and the product $\lambda_T \hat{n}_{\text{RNP}}^{\text{ss}} \approx 0.04 \ll 1$ places the system squarely in the non-depleted regime. Under these favorable conditions, molecular-scale optimizations can achieve their full potential.

Incorporating experimental values for our model system, $K_{D,T} = 1$ nM, $\eta = 1$, $\gamma_E = 1$, $k_{\text{cat},T} \sim 10^{-1}$, $k_{\text{cat,RNP}} \sim 10^{-5}$, $k_{\text{cat,apo}} \sim 10^{-5}$, $\pi c_s l^2 n_m \sim 3000$, $k_{\text{LoD}} \sim 100$, $r_0 = 4\mu$m, $n_m \sim 10^4/\mu\text{m}^2$, $k_{\text{off},T} \sim 1\text{s}^{-1}$, we estimate a detection limit in the 1-10 pM range, corresponding to approximately $10^5 - 10^6$ target copies per microliter. For targets near this detection threshold, the optimal measurement time approaches 80 minutes, while higher concentrations can be detected much more rapidly. Accordingly, this estimate and timing optimum apply to near-LoD operation and should not be treated as a universal optimum across all target-concentration regimes.

Our analysis suggests a hierarchical optimization strategy that addresses parameters according to their fundamental impact on performance. The first priority involves controlling macroscale depletion by ensuring $\lambda_T \hat{n}_{\mathrm{RNP}}^{\mathrm{ss}} < 1$ through appropriate bead parameter selection. Only after establishing this foundation do molecular-scale optimizations become effective. The second level focuses on nanoscale parameters. Surface density represents a powerful lever, with sensitivity improving as $n_m^{-3/4}$ in the non-depleted regime. Modern functionalization techniques can achieve densities approaching $5 \times 10^4$ molecules/$\mu$m$^2$, though steric crowding effects become significant at these levels. Guide-to-reporter ratios should be tailored to the specific background mechanisms, typically falling between 1-4 for most applications. Linker engineering deserves particular attention because combined linker length enters the sensitivity equation quadratically. PEG-based linkers in the 5-20 nm range typically provide optimal performance. The final optimization level involves operational parameters like measurement timing, temperature, and buffer composition. Time-resolved measurements can expand dynamic range by extracting concentration information from kinetic profiles rather than single timepoints. Temperature selection balances enhanced enzyme activity against reduced binding stability. Buffer optimization involves balancing ionic strength effects on binding affinity and enzyme activity while minimizing non-specific interactions. This multiscale optimization framework transforms the daunting complexity of diagnostic design into a manageable sequence of focused improvements. By systematically addressing depletion constraints, then molecular parameters, and finally operational conditions, developers can efficiently navigate toward optimal performance for their specific application requirements while avoiding common pitfalls that arise from piecemeal optimization approaches.

#### 4. Model Validation and Parameter Extraction from Experimental Data

Validating our theoretical framework against experimental data requires systematic parameter extraction that leverages different limiting behaviors. These fitting procedures correspond to the kinetic datasets shown in the main text (Fig. 2B-C), and the resulting parameters are used for the predictive sweeps shown in Fig. 3. We utilized kinetic fluorescence data collected under high and low target concentration regimes to extract the key model parameters through a sequential fitting approach. All experimental data, converted to MESF units, were normalized by the measured surface density $n_m = 1.7 \times 10^4$ molecules/$\mu$m$^2$ to enable direct comparison with our dimensionless theoretical expressions. The reactions were allowed to equilibrate based on the longer diffusion timescales for enzyme transport, ensuring that surface binding reached steady-state conditions before kinetic fluorescence measurements.

We began parameter extraction with high target concentration experiments where the initial reaction rates provide access to the intrinsic catalytic activity. Under these conditions, activated complexes are abundant and substrates are initially plentiful, minimizing accessibility constraints in the early stages of the reaction. Fitting the initial slopes of high concentration kinetic data yielded $\hat{k}_{\mathrm{cat},T} \approx 0.08$, or in dimensional form, $k_{\mathrm{cat},T} \approx 5 \times 10^{-6} \mu\mathrm{m}^2/\mathrm{s}$, representing the catalytic rate constant for surface-immobilized activated RNP complexes. Low target concentration experiments systematically probed the accessibility-limited regime where activated RNP complexes are sparse relative to available reporter molecules. Under these conditions, the limiting factor becomes how many substrate molecules each activated complex can effectively reach, making the substrate-side accessibility coefficient $c_s$ the dominant parameter. Initial fitting of linker length variations in the low target regime, using the previously determined catalytic rate constant, yielded $c_s \approx 20$.

However, this single-parameter accessibility model systematically overestimated the observed signal behavior in high target concentration experiments as the reactions progressed. The discrepancy arises because high target conditions initially create many activated complexes that progressively consume the available reporter molecules. As substrate depletion occurs during the reaction, the system transitions toward a substrate-limited regime reminiscent of classical Michaelis-Menten kinetics at high enzyme concentrations. In this depleted state, the limiting factor becomes how many activated complexes can reach each remaining substrate molecule, requiring introduction of the enzyme-side accessibility coefficient $c_e$. The complete bidirectional accessibility model recognizes that different experimental conditions probe distinct limiting behaviors. Low target experiments isolate $c_s$ by maintaining substrate-rich conditions with sparse activated complexes, while high target experiments reveal $c_e$ through the substrate depletion that occurs as reactions progress. This physical interpretation guided our fitting strategy: we used low target data to extract $c_s$, then employed the progression of high target concentration reactions to determine $c_e \approx 0.05$. The smaller value of $c_e$ suggests that activated complexes have lesser effective reach than the geometric accessibility of substrates to enzymes, likely due to differences in molecular size, conformational flexibility, and orientational constraints.

Background rate constants were extracted from dedicated control experiments designed to isolate non-specific catalytic activity. Apo-Cas only experiments, conducted with beads lacking guide RNAs but containing reporter RNA, directly measured the non-specific cleavage activity yielding $\hat{k}_{\text{cat,apo}} \approx 3 \times 10^{-5}$, or in dimensional terms, $k_{\text{cat,apo}} \approx 3 \times 10^{-4}\,\text{M}^{-1}\text{s}^{-1}$. Full no-target control experiments with surface-bound RNP complexes enabled extraction of the unactivated RNP background rate $\hat{k}_{\text{cat,RNP}} \approx 1.5 \times 10^{-5}$, or in dimensional terms, $k_{\text{cat,RNP}} \approx 10^{-9}\,\mu\text{m}^2\text{s}^{-1}$, many orders of magnitude lower than the target specific catalytic rate $k_{\text{cat},T}$. These background measurements are essential for accurate prediction of detection limits across different experimental conditions.

**Comparison with Solution-Phase Literature Values.** Our fitted parameter $k_{\text{cat},T} = 5 \times 10^{-6}\ \mu\text{m}^2/\text{s}$ converts to a molecular turnover number by multiplying by surface density: $k_{\text{cat}} = k_{\text{cat},T} \times n_m = 0.085\ \text{s}^{-1}$. The same Cas/guide/target/reporter system in bulk solution exhibits $k_{\text{cat}} \approx 30\ \text{s}^{-1}$, which aligns well with established CRISPR enzyme rates. Literature values for LbCas12a range from 0.07-17 $\text{s}^{-1}$[S1] and Cas13a systems span 17-740 $\text{s}^{-1}$ depending on substrates and conditions.[S2] Surface immobilization reduces our system's catalytic rate by nearly two orders of magnitude (from 30 to 0.085 $\text{s}^{-1}$), consistent with established effects of enzyme immobilization.[S3] Despite this substantial reduction in catalytic rate, the detection limits between surface and bulk systems differ by less than one order of magnitude. This suggests that the signal-to-background ratio scales differently than the absolute catalytic rate, with surface immobilization potentially providing geometric advantages that partially compensate for reduced enzymatic activity. These advantages include local concentration enhancement near tethered reporters and reduced effective interaction volume, which increase productive encounters despite lower intrinsic turnover.

#### 5. Quantification of tethered Cas13 reaction molecules on beads

To accurately quantify the number of Cas13 reaction molecules tethered to bead surfaces, we employed calibration beads (Quantum MESF Beads, Bangs Laboratories) that facilitate approximate estimation of number of surface-bound fluorophores.[S4] These calibration beads are surface-labelled with defined quantities of fluorophores across five distinct bead populations, enabling the generation of a precise fluorescence standard curve.[S5]

In our experimental protocol, AlexaFluor 647-labelled guide RNA and FITC-labelled reporter molecules were added to the beads in solution at varying molar ratios. Figure S1A presents the quantified surface densities of guide RNA and reporter molecules in units of Molecules of Equivalent Soluble Fluorophores (MESF) across varying solution ratios. Due to differential binding affinities and surface saturation effects, the ratios of guide RNA to reporter molecules tethered to the bead surface differed from their initial solution ratios as shown in Figure S1B. Excess unbound guide RNA and reporter molecules were removed via thorough washing steps.

To quantify the tethered molecules, beads with surface-bound AlexaFluor 647-labelled guide RNA and FITC-labelled reporters were imaged using an epifluorescence microscope under identical conditions alongside the calibration beads. Imaging was performed concurrently to minimize variability. The fluorescence intensity obtained from the calibration beads established a robust standard curve, correlating fluorescence intensity to the precise number of fluorophores per bead.

This calibration method allowed accurate quantification and precise control over the number of surface-tethered guide RNA and reporter molecules. The resulting quantified data was then integrated into our analytical framework, enabling prediction of critical SURFCas assay parameters for each experimental condition.

#### 6. Experimental optimization of SURFCas reaction parameters

**Combination of Multiple Distinct Guide RNAs on beads.** To enhance the sensitivity of the SURFCas assay, we tested the efficacy of tethering multiple distinct guide RNAs targeting separate regions of the same target RNA on a single bead surface. During the Cas13 reaction, target RNA molecules bound to ternary Cas13 complexes undergo cis-cleavage, generating RNA fragments. These fragments can subsequently engage other distinct Cas13-guide RNA complexes tethered to the bead surface. Such sequential engagement amplifies Cas13 activation, especially in assays without prior RNA amplification steps where target RNA concentration is inherently limiting.[S6]

We assessed this approach using three distinct guide RNAs: crRNA 2, crRNA 4, and crRNA D7, each targeting separate regions of the SARS-CoV-2 genome. We compared individual guide RNAs against a combined condition with all three guide RNAs tethered on the same bead. In the combined condition, the total number of guide RNA molecules per bead was consistent across experiments but evenly distributed among the three crRNAs. Synthetic RNA targets, serving as proxies for full-length SARS-CoV-2 RNA, were added at concentrations of 1e6 and 5e5 copies/$\mu$L. Our results demonstrated that combining multiple guide RNAs substantially improved Cas13 activation, reducing the assay's limit of detection (LOD) from 1e6 copies/$\mu$L for individual guide RNAs to 5e5 copies/$\mu$L for the combined guide RNA condition (Figure S2).

**Impact of Guide RNA Tethering Orientation.** The orientation of guide RNA tethering significantly impacts Cas13 reaction efficiency. We compared tethering at the 5' end versus the 3' end of guide RNA molecules. Our results indicated a pronounced increase in reaction rate and sensitivity when guide RNAs were tethered at the 3' end compared to the 5' end. Specifically, tethering at the 3' end enhanced the LOD by an order of magnitude—from 1e7 copies/$\mu$L (5' tethering) to 1e6 copies/$\mu$L (3' tethering), as shown in Figure S3.

**Influence of PEG on Reaction Performance.** We investigated the role of molecular crowding by varying concentrations of polyethylene glycol (PEG-8000) in the reaction mixture. At high target RNA concentrations (1e8 copies/$\mu$L), we observed similar endpoint fluorescence across a range of PEG concentrations (0–15 %), suggesting that the reaction kinetics were not diffusion-limited under abundant RNA conditions (Figure S4A). However, at RNA concentrations near the LOD (1e6 copies/$\mu$L), the optimal performance was achieved with 7.5 % PEG-8000 (Figure S4B). Higher concentrations of PEG-8000 (e.g., 15 %) significantly decreased reaction efficiency. This effect can be explained by PEG-induced macromolecular crowding, which initially raises the effective concentration of target RNA near the bead surface, accelerating target binding and collateral cleavage.[S7] Nevertheless, excessive PEG concentrations lead to increased viscosity and reduced water activity, ultimately impairing molecular diffusion and offsetting the benefits of molecular crowding.

#### Supplementary Figures

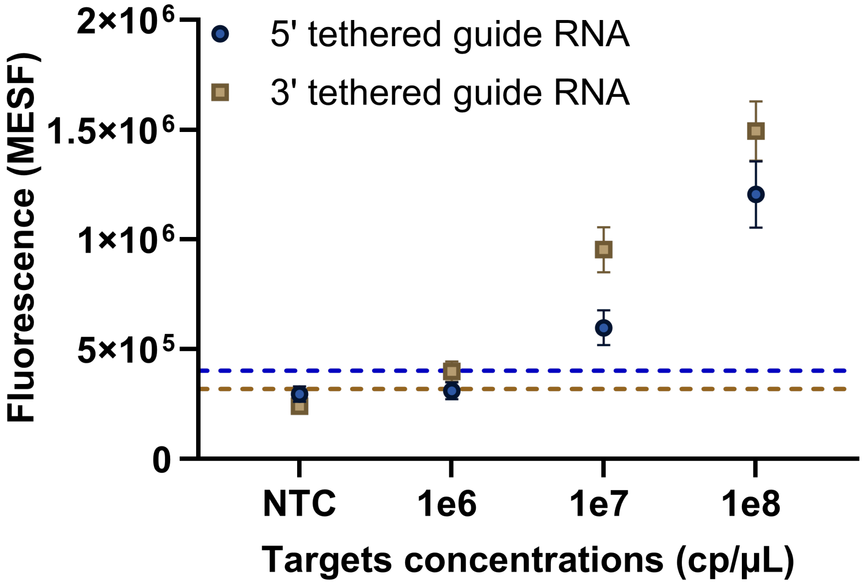


Figure S1: **Effect of guide RNA tethering orientation on SURFCas performance** Comparison of reaction performance across varying target RNA concentrations and limit of detection (LOD) when guide RNAs are tethered at the 5' end versus the 3' end. Dashed lines indicate the threshold for determining LOD, defined as the mean of the no-template control (NTC) condition plus three times its standard deviation (mean NTC + 3 × SD).

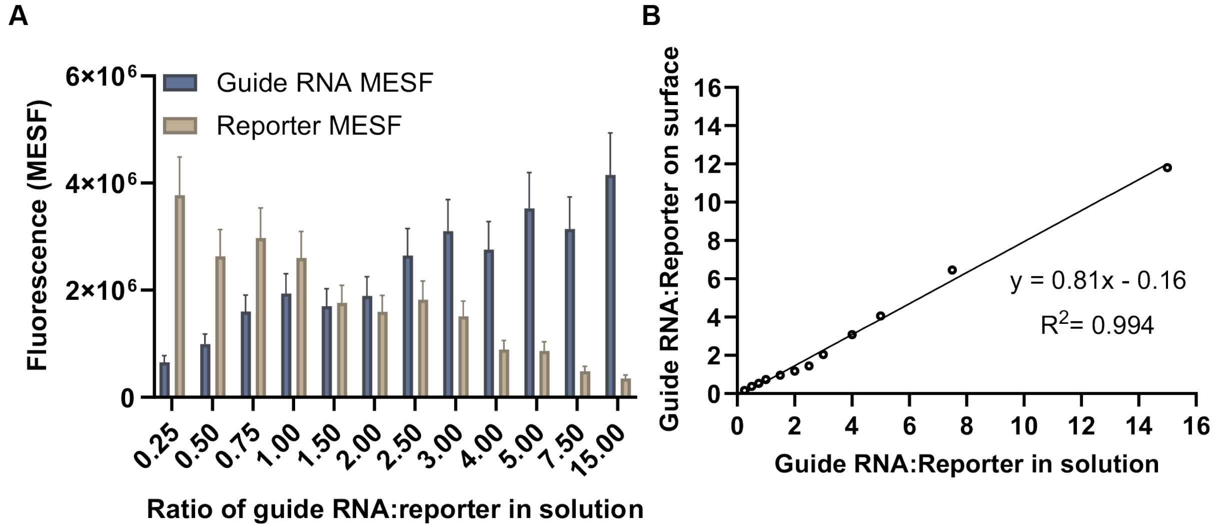


Figure S2: **Quantification of guide RNA and reporter molecules tethered on beads.** (A) MESF of labelled guide RNA and reporter molecules bound to bead surfaces after incubation in varying solution ratios. Error bars represent the standard deviation of bead fluorescence intensities across the total number of beads per condition. (B) Relationship between the ratio of guide RNA to reporter molecules added in solution and their resulting ratio on bead surfaces.

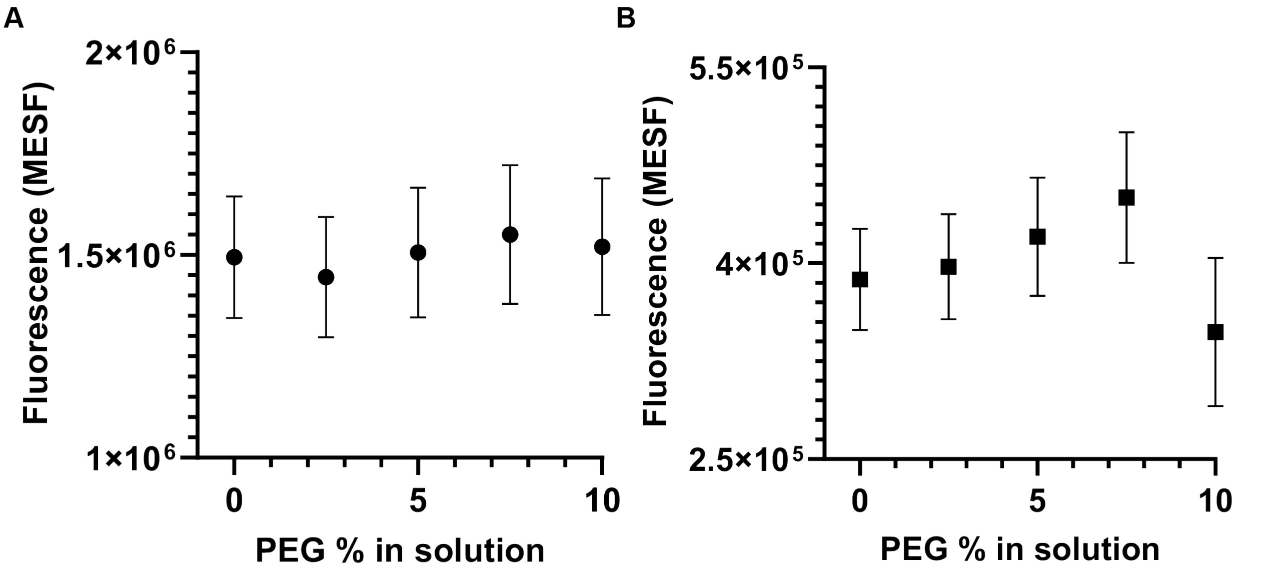


Figure S3: **Effect of adding soluble PEG in reaction mixture** (A) Influence of adding different concentrations of soluble PEG-8000 at target RNA concentration of 1e8 copies/$\mu$L (B) Influence of adding different concentrations of soluble PEG-8000 at target RNA concentration of 1e6 copies/$\mu$L. Error bars represent the standard deviation of bead fluorescence intensities across the total number of beads per condition.

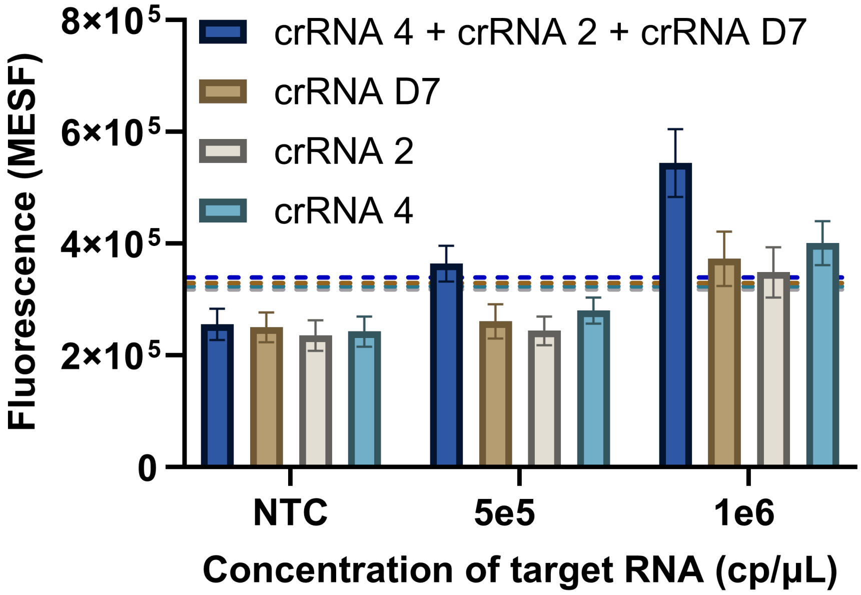


Figure S4: **Effect of tethering multiple distinct guide RNAs for the same target RNA molecule per bead** Comparison of Cas13 reaction performance across varying target RNA concentrations for three different guide RNAs tethered to bead exclusively or in combination. Dashed lines indicate the threshold for determining LOD, defined as the mean of the no-template control (NTC) condition plus three times its standard deviation (mean NTC + 3 × SD).

### Supplementary Table

Table S1: **Oligonucleotides used in this study.** Sequences of crRNAs, reporters, and target RNAs.

| Oligonucleotide description | Sequence |
|---|---|
| crRNA 4 targeting SARS-CoV-2 genome | rGrArCrCrArCrCrCrCrArArArArArUrGrArArGrGrGrGrArCrUrArArArArCrUrUrUrGrCrGrGrCrCrArArUrGrUrUrUrGrUrArA |
| crRNA 2 targeting SARS-CoV-2 genome | rGrArCrCrArCrCrCrCrArArArArArUrGrArArGrGrGrGrArCrUrArArArArCrGrGrUrCrCrArCrCrArArArCrGrUrArArUrGrCrG |
| crRNA D7 targeting SARS-CoV-2 genome | rGrArCrCrArCrCrCrCrArArArArArUrGrArArGrGrGrGrArCrUrArArArArCrCrArCrArGrUrCrArUrArArUrCrUrArUrGrUrUrA |
| Reporter | /56-FAM/rUrUrUrUrU/3IABkFQ/ |
| 5' surface tethered crRNA 4 with 5 PEG spacer molecules (linker length of ∼12.6 nm) (for 5' vs 3' data) | /5BiotinTEG//iSp18//iSp18//iSp18//iSp18//iSp18/rGrArCrCrArCrCrCrCrArArArArArUrGrArArGrGrGrGrArCrUrArArArArCrUrUrUrGrCrGrGrCrCrArArUrGrUrUrUrGrUrArA |
| 3' surface tethered crRNA 4 with no spacer | rGrArCrCrArCrCrCrCrArArArArArUrGrArArGrGrGrGrArCrUrArArArArCrUrUrUrGrCrGrGrCrCrArArUrGrUrUrUrGrUrArA/3BioTEG/ |
| 3' surface tethered crRNA 4 with 5 PEG spacer molecules (linker length of ∼12.6 nm) | rGrArCrCrArCrCrCrCrArArArArArUrGrArArGrGrGrGrArCrUrArArArArCrUrUrUrGrCrGrGrCrCrArArUrGrUrUrUrGrUrArA/iSp18//iSp18//iSp18//iSp18//iSp18//3BioTEG/ |
| 3' surface tethered crRNA 4 with 20 PEG spacer molecules (linker length of ∼50.4 nm) | rGrArCrCrArCrCrCrCrArArArArArUrGrArArGrGrGrGrArCrUrArArArArCrUrUrUrGrCrGrGrCrCrArArUrGrUrUrUrGrUrArA/iSp18//iSp18//iSp18//iSp18//iSp18//iSp18//iSp18//iSp18//iSp18//iSp18//iSp18//iSp18//iSp18//iSp18//iSp18//iSp18//iSp18//iSp18//iSp18//iSp18//3BioTEG/ |
| 3' surface tethered photoactivable crRNA 4 with 20 PEG spacer molecules (linker length of ∼50.4 nm) | TTTTTTTTTTTTTTTTTTT/iSpPC/CrGrArCrCrArCrCrCrCrArArArArArUrGrArArGrGrGrGrArCrUrArArArArCrUrUrUrGrCrGrGrCrCrArArUrGrUrUrUrGrUrArA/iSp18//iSp18//iSp18//iSp18//iSp18//iSp18//iSp18//iSp18//iSp18//iSp18//iSp18//iSp18//iSp18//iSp18//iSp18//iSp18//iSp18//iSp18//iSp18//iSp18//3BioTEG/ |
| 3' surface tethered crRNA 4 with AlexaFluor 647 label | /5Alex647N/rGrArCrCrArCrCrCrCrArArArArArUrGrArArGrGrGrGrArCrUrArArArArCrUrUrUrGrCrGrGrCrCrArArUrGrUrUrUrGrUrArA/iSp18//iSp18//iSp18//iSp18//iSp18//iSp18//iSp18//iSp18//iSp18//iSp18//iSp18//iSp18//iSp18//iSp18//iSp18//iSp18//iSp18//iSp18//iSp18//iSp18//3BioTEG/ |
| 3' surface tethered crRNA 2 with 5 PEG spacer molecules (linker length of ∼12.6 nm) | rGrArCrCArCrCrCrCArArArArArUrGrArArGrGrGrGrArCrUrArArArArCrGrGrUrCrCrArCrCrArArACrGrUrArArUrGrCrG/iSp18//iSp18//iSp18//iSp18//iSp18//3BioTEG/ |
| 3' surface tethered crRNA D7 with 5 PEG spacer molecules (linker length of ∼12.6 nm) | rGrArCrCrArCrCrCrCrArArArArArUrGrArArGrGrGrGrArCrUrArArArArCrCrArCrArGrUrCrArUrArArUrCrUrArUrGrUrUrA/iSp18//iSp18//iSp18//iSp18//iSp18//3BioTEG/ |
| 5' surface tethered reporter with no spacer | /56-FAM//iBiodT/rUrUrUrUrU/3IABkFQ/ |
| 5' surface tethered reporter 2 with 5 PEG spacer molecules (linker length of ∼12.6 nm) | /5BiotinTEG//iSp18//iSp18//iSp18//iSp18//iSp18//iFluorT/rUrUrUrUrU/3IABkFQ/ |
| 5' surface tethered reporter with 20 PEG spacer molecules (linker length of ∼50.4 nm) | /5BiotinTEG//iSp18//iSp18//iSp18//iSp18//iSp18//iSp18//iSp18//iSp18//iSp18//iSp18//iSp18//iSp18//iSp18//iSp18//iSp18//iSp18//iSp18//iSp18//iSp18//iSp18//iFluorT/rUrUrUrUrU/3IABkFQ/ |
| Synthetic target RNA for crRNA 4 | rGrGrArArCrUrGrArUrUrArCrArArArCrArUrUrGrGrCrCrGrCrArArArUrUrGrCrArCrArA |
| Synthetic target RNA for crRNA 2 | rUrGrCrArCrCrCrCrGrCrArUrUrArCrGrUrUrUrGrGrUrGrGrArCrCrCrUrCrArGrArUrU |
| Synthetic target RNA for crRNA D7 | rGrUrUrGrGrUrUrUrUrArArCrArUrArGrArUrUrArUrGrArCrUrGrUrGrUrCrUrCrUrUrU |
